# Phylodynamic signatures in the emergence of community-associated MRSA

**DOI:** 10.1101/2021.04.30.442212

**Authors:** Eike Steinig, Izzard Aglua, Sebastián Duchêne, Michael T. Meehan, Mition Yoannes, Cadhla Firth, Jan Jaworski, Jimmy Drekore, Bohu Urakoko, Harry Poka, Clive Wurr, Eri Ebos, David Nangen, Elke Müller, Peter Mulvey, Charlene Jackson, Anita Blomfeldt, Hege Vangstein Aamot, Moses Laman, Laurens Manning, Megan Earls, David C. Coleman, Andrew Greenhill, Rebecca Ford, Marc Stegger, Muhammed Ali Syed, Bushra Jamil, Stefan Monecke, Ralf Ehricht, Simon Smith, William Pomat, Paul Horwood, Steven Y.C. Tong, Emma McBryde

## Abstract

Community-associated, methicillin-resistant *Staphylococcus aureus* (MRSA) lineages have emerged in many geographically distinct regions around the world during the past 30 years. Here, we apply consistent phylodynamic methods across multiple community-associated MRSA lineages to describe and contrast their patterns of emergence and dissemination. We generated whole genome sequencing data for the Australian sequence type (ST) 93-MRSA-IV from remote communities in Far North Queensland and Papua New Guinea, and the Bengal Bay ST772-MRSA-V clone from metropolitan communities in Pakistan. Increases in the effective reproduction number (R_e_) and sustained transmission (R_e_ > 1) coincided with spread of progenitor methicillin-susceptible *S. aureus* (MSSA) in remote northern Australia, dissemination of the ST93-MRSA-IV genotype into population centers on the Australian East Coast, and subsequent importation into the highlands of Papua New Guinea and Far North Queensland. Analysis of a ST772-MRSA-V cluster in Pakistan suggests that sustained transmission in the community following importation of resistant genotypes may be more common than previously thought. Applying the same phylodynamic methods to existing lineage datasets, we identified common signatures of epidemic growth in the emergence and epidemiological trajectory of community-associated *S. aureus* lineages from America, Asia, Australasia and Europe. Surges in R_e_ were observed at the divergence of antibiotic resistant strains, coinciding with their establishment in regional population centers. Epidemic growth was also observed amongst drug-resistant MSSA clades in Africa and northern Australia. Our data suggest that the emergence of community-associated MRSA and MSSA lineages in the late 20th century was driven by a combination of antibiotic resistant genotypes and host epidemiology, leading to abrupt changes in lineage-wide transmission dynamics and sustained transmission in regional population centers.

---

***S****taphylococcus aureus* is an opportunistic pathogen that causes a variety of clinical manifestations, from superficial skin and soft tissue infections to life-threatening systemic diseases, including bloodstream infections and necrotizing pneumonia (1). Unfortunately, treatment has been complicated by the rapid emergence of antibiotic resistance worldwide. In the last few decades, a series of distinct *S. aureus* lineages, defined by multilocus sequence types (ST), have emerged in healthcare, community and agricultural settings around the world (2–4). Strains of methicillin-resistant *S. aureus* (MRSA) within some of these lineages have traditionally disseminated in hospitals, where acquisition of mutations and mobile genetic elements - such as the staphylococcal cassette chromosome mec (SCC*mec*) - promote persistence under high antibiotic selection pressure (5, 6). Since the 1990s, however, antibiotic-resistant community-associated clones without epidemiological links to hospitals have emerged around the world, subsequently replacing other regionally prevailing lineages (2). Community-associated MRSA strains tend to be virulent, infect otherwise healthy people, and are frequently exported from the regions in which they emerged (6). While considered less resistant to antibiotics than healthcare-associated strains, evidence from multiple global and regional whole-genome datasets suggests that their emergence is associated with the acquisition of specific resistance mutations and mobile elements (7–15).

Epidemiological and genomic evidence for historical and ongoing circulation of MSSA progenitor populations exists for nearly all community-associated lineages of interest (7–15). Strong contemporary evidence comes from the Australian ST93 lineage, whose ancestral MSSA strains continue to circulate amongst remote communities in the Northern Territory (10, 16). In addition, a symplesiomorphic clade of ST8-MSSA progenitor strains has been found circulating in Africa (7), having diverged prior to the emergence of the ancestral ST8-MSSA in Europe during the 19th century, which then spread to the Americas where it diverged into the ST8-USA300 (MRSA) sublineages in the 20th century (13, 14). Local circulation of progenitor MSSA strains in Romania is documented for the European ST1-MRSA sublineage (11, 17, 18). Emergence of ST80-MRSA in Europe has epidemiological connections to North West Africa through importation of MSSA cases in French legionaires (8, 19). While few ancestral strains have been sampled, ST772-MRSA-V is thought to have emerged from local MSSA populations in the Bengal Bay area, with the first isolates from 2004 collected in Bangladesh and India, coinciding with the rise of a multidrug-resistant MRSA clade on the Indian subcontinent (15, 20). Even less is known about the origins of the ST59 clone, which produced an MRSA epidemic in Taiwan, but had previously diverged into a (largely) MSSA sister clade in the United States (9).

Subsequent global dissemination of emergent MRSA clades has frequently been linked to travel and family history in their source region (7–15). For example, nearly 60% of isolates included from a global study on the dissemination of the ST772-MRSA-V clone had family contacts or travel history on the Indian subcontinent (15). However, to date, community strains tend to cause small-scale outbreaks, consisting of local transmission chains and household clusters failing to become endemic in the community (7, 10, 15, 21–23). Some notable exceptions include several USA300 clades (ST8-MRSA-IV genotype) in Colombia, Gabon and France, as well as the Australian (ST93-MRSA-IV genotype) featuring a transmission event into the M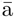ori and Pacific Islander in metropolitan Auckland, New Zealand (NZ) (7, 10, 13). Additional evidence for successful recruitment arises from molecular surveillance of ST80 and ST1, as well as from genomic surveillance of ST152-MSSA in the Middle East (8, 12, 18, 24).

The distinct regional distribution of community-associated lineages is observed in stark contrast to healthcare-associated strains, that tend to spread rapidly in local healthcare systems, often following international dissemination (5, 25, 26). The evolutionary and epidemiological trajectory of community-associated lineages is currently not known, although indications are that the prevalence of some lineages and sublineages have declined over the decade, including the North American USA300 clade (27). However, in disadvantaged and remote communities, such as in northern Australia (28), community-associated MRSA rates are some of the highest in the country and have been increasing by around 4% per year, contrary to all other healthcare jurisdictions in Australia over the same period of time (29, 30). In addition, extremely remote populations in the Pacific Islands, such as in Papua New Guinea, which borders Australia in the Torres Strait, have reported outbreaks of community-associated osteomyelitis infections caused by *S. aureus*, but little is known about the extent of these outbreaks (31, 32).

While these data have contributed to a deeper understanding of community-associated lineage emergence, questions remain about the drivers behind these seemingly convergent events in the late 20th century. Increases in the effective population size (N_e_) have been observed in some lineages, coinciding with the acquistion of antibiotic resistance but these analyses have not been conducted in for all relevant sequence types (8, 9, 11, 12). While historical and contemporary data on MSSA progenitor populations is limited in most lineages, we note that these populations tend to be geographically distinct, and that emergence of resistant genotypes occurs rapidly in industrialized host populations, such as on the Indian subcontinent (ST772), the Australian East Coast (ST93), in central Europe (ST1, ST80) and North America (ST8). In addition, it is not clear whether sustained transmission — characterized by an effective reproduction number (R_e_) exceeding a threshold value of one, and remaining above that threshold for a period of time — has occurred following the emergence and transmission of community-associated strains, and whether drug-resistant strains are capable of becoming endemic following their exportation. Bayesian phylodynamic methods have been extensively used in viral epidemics to infer key epidemiological parameters and changes in transmission dynamics (Δ R_e_) from phylogenetic trees (33–36), allowing for simultaneous assessment of genomic and epidemiological changes in emerging pathogens. However, phylodynamic applications have been limited for bacterial datasets due to their relatively more complex genome evolution, lack of meta-data and sufficient longitudinal isolate collections (37, 38). Virtually nothing is known about the transmission dynamics of *S. aureus*. Synthesizing the available evidence, we hypothesize that interactions between genomic and epidemiological factors create the conditions necessary for sustained transmission in the local environment, and have contributed to the emergence of community-associated MRSA.

Here, we investigate the genome evolution and transmission dynamics of emergent community-associated MRSA lineages using comparative phylodynamic methods to describe and contrast patterns of emergence and spread. We first examine the genomic epidemiology and transmission dynamics of community-associated *S. aureus* from the remote highlands of Papua New Guinea (PNG) and communities in northern Australia (Far North Queensland, FNQ). Using additional samples of the Bengal Bay clone (ST772) from Pakistan (39) and global lineage-resolved sequence data (7–15), we discover signatures in the effetive reproduction number that suggest a combination of resistance acquistion and epidemic growth in populations centers as key drivers in the emergence of community-associated MRSA.

## Results

We sequenced 187 putative *S. aureus* isolates from remote PNG (2012 - 2018) and FNQ (Torres and Cape / Cairns and Hinterland jurisdictions, 2019) using Illumina short-reads (Fig. 1, Supplementary Tables). Genotyping identified the Australian MRSA clone (ST93-MRSA-IV) as the main cause of paediatric osteomyelitis (31) in the highland towns of Kundiawa and Goroka (n_Kundiawa_ = 33/42, n_Goroka_ = 30/35). The remaining isolates from osteomyelitis cases in Kundiawa and Goroka belonged to an assortment of sequence types (ST5, ST25, ST88), single locus variants of ST1247 (n = 1) and ST93 (n = 2), coagulase-negative staphylococci including *S. lugdunensis* (n = 1), *S. delphini* (n = 1) and *Mammaliicoccus sciuri* (n = 1), as well as a neonatal hospital cluster of invasive ST243 (clonal complex 30, n = 9) (Fig. 1A). FNQ isolates sampled in 2019 were largely identified as ST93-MRSA-IV (n_FNQ_ = 68/91) on a background of various other lineages, including one infection with *S. argenteus* (Fig. 1A, Supplementary Tables).

**Fig. 1.**
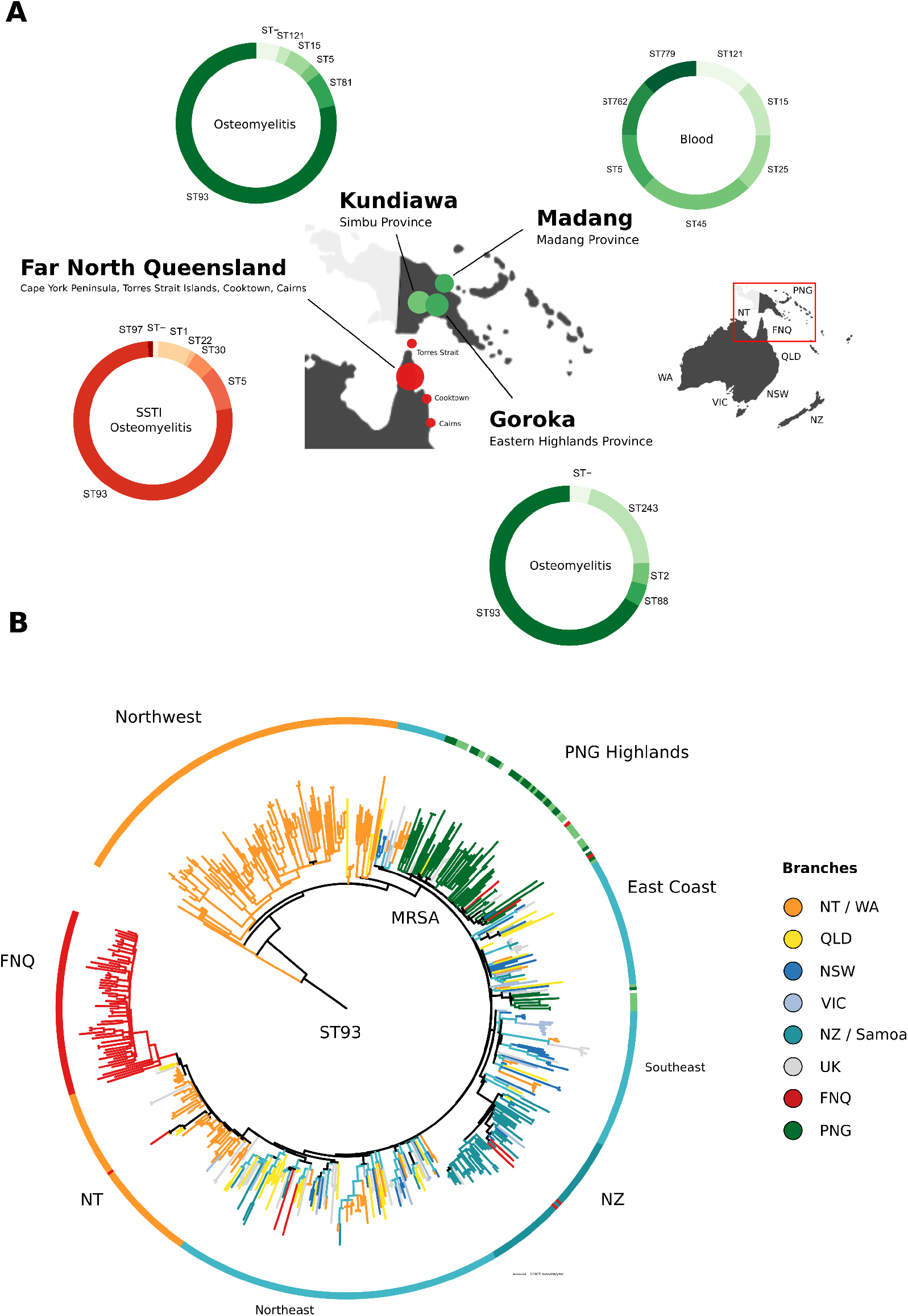
Genomic epidemiology of *Staphylococcus aureus* outbreak isolates from Papua New Guinea (n = 95) and Far North Queensland (n = 89). (**A**) Map of sampling locations, multilocus sequence types and predominant symptoms of patients (ring annotation) (**B**) Global evolutionary history of the Australian lineage (ST93) showing the rooted maximum-likelihood phylogeny constructed from a non-recombinant core-genome SNP alignment (n = 575) and major regional geographical structure in the evolutionary history of the clone (branch colors). ST93 emerged in remote communities of North West Australia and acquired SCC*mec*-IV, spreading to the Australian East Coast (blues, yellow), remote northern Australian communities (orange, red), the remote highlands of Papua New Guinea (green) and into Auckland communities in New Zealand (seagreen).

ST93-MRSA-IV strains from PNG and FNQ were contextualised within the global sequence diversity of the lineage (ST93, n = 444) to determine strain provenance using a maximum-likelihood (ML) phylogeny constructed from non-recombinant core-genome SNPs (Fig. 1B, n = 575, 6648 SNPs). The resulting tree topology recapitulated our previous analysis of the lineage, confirming its origin from extant MSSA strains circulating in remote Indigenous communities of north-western Australia (10). The main divergence event of ST93-MRSA on the Australian East Coast (AEC) coincided with acquisition of SCC*mec*-IV (Fig. 1B). Isolates from PNG formed a major (n = 55) and minor (n = 8) clade in the ML tree, consisting of mixed strains from Goroka and Kundiawa (Fig. 1A, 1B, green). The major clade contained sporadic isolates sampled in Queensland, FNQ, and New South Wales (n = 3) indicating regional transmission from PNG (Fig. 1B). The FNQ cluster derived from a Northern Territory (NT) clade that itself appears to have been a re-introduction of ST93-MRSA-IV from Austalia’s East Coast into the Northern Territory (Fig. 1B). Sporadic isolates sampled in FNQ were imported from other locations, including the North Eastern ST93-MRSA-IV circulation, the NT, as well as NZ and PNG (red branches outside of FNQ cluster in Fig. 1B). Sporadic transmission into FNQ most likely occurred through Cairns, which is the regional hub of the area, has an international airport and is frequented by visitors from the region.

### Regional transmission dynamics of the Australian community clone (ST93-MRSA-IV)

We next used fast maximum-likelihood methods (40, 41) (Fig. S1, Fig. S2) as well as Bayesian coalescent skyline (42) and birth-death skyline (33) models for serially (PNG) and contemporaneously sampled isolates (FNQ) in `BEAST2` (34) to infer time-scaled phylogenies and estimate epidemiological parameters for the ST93-MRSA-IV clone, including changes in R_e_ and effective population size (N_e_) over time (Fig. 2A, Table 1). Previous genomic studies have noted increases in N_e_ in the emergence of several community lineages (8, 9, 11, 12) but data was not available for all lineages (7, 10, 20), and no studies had previously used birth-death skyline models to investigate changes in R_e_. Lineage-wide transmission dynamics of the Australian clone ST93 indicate successive surges in R_e_ at the divergence of extant MSSA strains in the Northern Territory (NT), at acquisition of SCC*mec*-IVa and spread on the AEC, and upon recruitment into PNG, NZ and FNQ communities (Fig. 2A). The clone became epidemic (R_e_ > 1) soon after the emergence of an extant MSSA clade in the NT (MRCA = 1990, 95% credible interval, CI: 1988 - 1992), coinciding with the first sample (1991) from the NT in our retrospective collection (Fig. 2A). When the clone was first described in southern Queensland in 2000 (20) a resistant clade ST93-MRSA-IV had just established transmission in East Coast population centers (QLD, NSW, VIC) following the acquisition of SCC*mec*-IV around 1994 (95% CI: 1993 - 1995) (Fig. 1B, Fig. 2C). We estimate that the introduction of ST93-MRSA-IV into PNG occurred in the early 2000s (MRCA = 2000, 95% CI: 1998 – 2003, Fig. 2A) soon after establishment on the East Coast of Australia. In contrast, introduction of ST93-MRSA-IV into FNQ occurred more recently (MRCA = 2007, 95% CI: 2005 - 2009).

**Table 1.** Birth-death skyline median posterior estimates for global community-associated *S. aureus* lineages and sublineages.

| Lineage | n | n<sub>MSSA</sub> | MRCA | 95% CI | R<sub>e</sub> | 95% CI | Clock rate* | Notes | References |
| --- | --- | --- | --- | --- | --- | --- | --- | --- | --- |
| **Sequence types** |  |  |  |  |  |  |  |  |  |
| ST93 | 575 | 109 | 1983 | 1982 - 1983 | Δ | Figs. 2, 4 | $3.18 \times 10^{-04}$ | Australia | this study, (10, 16) |
| ST772 | 359 | 36 | 1970 | 1965 - 1975 | Δ | Figs. 3, 4 | $4.11 \times 10^{-04}$ | South Asia | this study, (15) |
| ST80 | 215 | 23 | 1977 | 1974 - 1979 | Δ | Fig. 4 | $5.93 \times 10^{-04}$ | Europe | (8, 19) |
| ST8 | 207 | 64 | 1860 | 1849 - 1871 | Δ | Fig. 4 | $1.95 \times 10^{-04}$ | Americas | (7, 13) |
| ST1 | 190 | 10 | 1987 | 1985 - 1989 | Δ | Fig. 4 | $4.26 \times 10^{-04}$ | Europe | (11, 17) |
| ST59 | 154 | 32 | 1958 | 1952 - 1964 | Δ | Fig. 4 | $3.69 \times 10^{-04}$ | US-Taiwan | (9) |
| ST152 | 117 | 61 | 1958 | 1947 - 1962 | Δ | Fig. 4 | $4.79 \times 10^{-04}$ | Europe | (12) |
| **MRSA sublineages** |  |  |  |  |  |  |  |  |  |
| ST93-MRSA-EastCoast | 278 | 2 | 1994 | 1993 - 1995 | 1.57 | 1.11 - 2.29 | $3.18 \times 10^{-04}$ | | (10) |
| ST93-MRSA-FNQ | 61 | 0 | 2007 | 2005 - 2009 | 1.55 | 1.08 - 2.27 | $3.18 \times 10^{-04}$ | | this study, (29) |
| ST93-MRSA-PNG | 65 | 3 | 2000 | 1998 - 2003 | 1.61 | 1.13 - 2.40 | $3.18 \times 10^{-04}$ | | this study, (31) |
| ST93-MRSA-NT | 62 | 1 | 1999 | 1998 - 2001 | 0.97 | 0.72 - 1.25 | $3.18 \times 10^{-04}$ | | (10, 16) |
| ST93-MRSA-NZ | 51 | 1 | 2002 | 2001 - 2004 | 1.09 | 0.79 - 1.48 | $3.18 \times 10^{-04}$ | | (10, 16) |
| ST772-MRSA-Pakistan | 25 | 0 | 2002 | 2000 - 2003 | 1.36 | 1.08 - 1.79 | $4.11 \times 10^{-04}$ | | this study, (39) |
| ST152-MRSA-Europe | 53 | 0 | 1989 | 1986 - 1991 | 1.47 | 1.07 - 2.12 | $4.79 \times 10^{-04}$ | | (12) |
| ST8-USA300-NorthAmerica | 73 | 3 | 1983 | 1979 - 1986 | 1.60 | 1.13 - 2.36 | $1.95 \times 10^{-04}$ | ACME** | (7, 13) |
| ST8-USA300-SouthAmerica | 32 | 0 | 1978 | 1973 - 1982 | 1.56 | 1.10 - 2.31 | $1.95 \times 10^{-04}$ | COMER** | (7, 13) |
| ST8-USA300-Gabon | 17 | 0 | 1993 | 1991 - 1995 | 1.58 | 1.11 - 2.34 | $1.95 \times 10^{-04}$ | | (7) |
| ST59-MRSA-Taiwan | 87 | 2 | 1979 | 1976 - 1981 | 1.46 | 1.07 - 2.11 | $3.69 \times 10^{-04}$ | | (9) |
| **MSSA sublineages** |  |  |  |  |  |  |  |  |  |
| ST93-MSSA-Australia | 96 | 84 | 1990 | 1988 - 1992 | 1.54 | 1.09 - 2.25 | $3.18 \times 10^{-04}$ | *blaZ, ermC* | (10) |
| ST8-MSSA-WestAfrica | 23 | 21 | 1979 | 1975 - 1982 | 1.54 | 1.09 - 2.26 | $1.95 \times 10^{-04}$ | *blaZ, dfrG, tetK, fosD* | (7) |
| ST152-MSSA-WestAfrica | 42 | 40 | 1975 | 1972 - 1978 | 1.55 | 1.10 - 2.25 | $4.79 \times 10^{-04}$ | *blaZ* | (12) |

\* sublineage clock rates fixed at lineage-wide estimated median, Δ changes in R<sub>e</sub> over time (Figs. 2, 3 4), \*\* COMER - copper and mercury resistance element; ACME - arginine catabolic mobile element

**Fig. 2.**
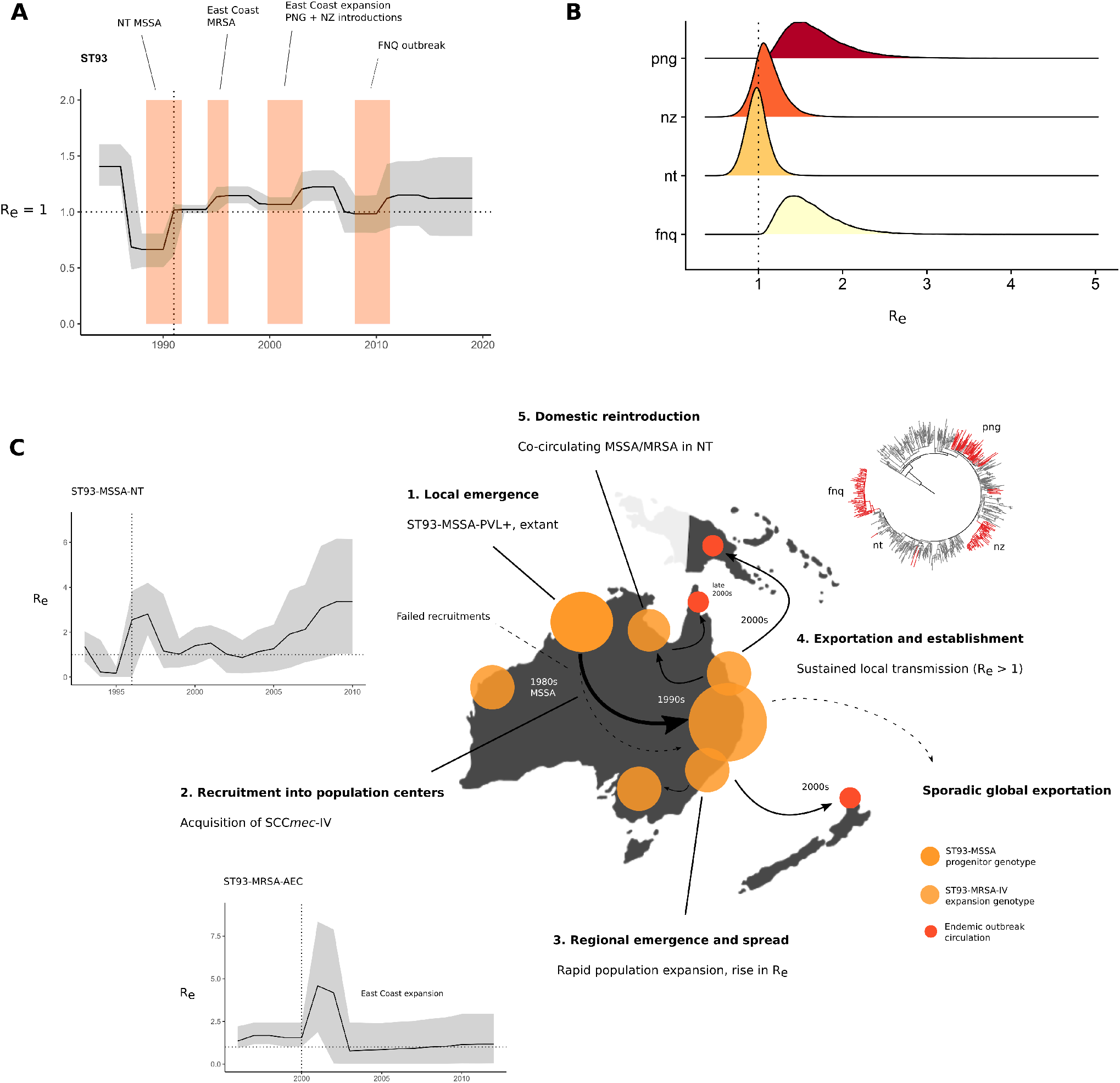
Phylodynamic signatures and parameter estimates for the Queensland clone (ST93) (**A**) Changes in the effective reproduction number (R_e_) over time, showing the 95% credible interval (CI) intervals of the MRCA of clade divergence events of ST93-MSSA and -MRSA (colors) using the birth-death skyline model (**B**) R_e_ posterior density distributions for introductions in Papua New Guinea (PNG), Far North Queensland (FNQ), New Zealand (NZ) and re-introduction of the MRSA genotype to the Northern Territory (NT) (**C**) Events in the emergence and regional dissemination of the Queensland clone, with maximum-likelihood phylogenies and branch colors indicating major sub-clades and divergence events in the emergence of ST93. Vertical lines in skyline plots indicate the year of first sample from the lineage or clade, horizontal lines indicate the epidemic threshold of R_e_ = 1. (**1**) Local emergence of ST93-MSS in remote Indigenous communities of North Western Australia. Some sporadic transmission to QLD occured in this clade. ST93-MSSA continue to circulate in the Northern Territory and R_e_ > 1 estimates indicate that strains continue to spread (inset plot). (**2**) When the lineage acquired SCC*mec*-IV it spread to East Australian coastal states (Queensland, Victoria, New South Wales) coinciding with a population growth and increase in transmission with a spike at initial recruitment (inset plot) (**3**) ST93-MRSA-IV continues to sprad in the eastern coastal states (QLD, NSW, VIC) (4) From the globally connected East Coast population centers, ST93-MRA-IV spread overseas (particularly to the U.K.) but also establishes sustained transmission in remote Far North Queensland and regionally in the highlands of Papua New Guinea. (5) The Far North Queensland outbreak is derived from a ST93-MRA-IV clade in the Northern Territory co-circulating with the ongoing ST772-MSSA epidemic.

Birth-death skyline models with fixed lineage-wide substitution rates were additionally applied to regional sublineages and -clades of ST93 (Methods), including the introductions into PNG, FNQ, New Zealand (NZ), and the re-introduction into the Northern Territory (Fig. 2B, Table 1: sublineages). We observed sustained transmission in PNG (R_e_ = 1.61, 95% CI: 1.13 – 2.40) and FNQ (R_e_ = 1.55, 95% CI: 1.08 – 2.44). Sustained transmission may have occurred in the Northern Territory re-introduction of the MRSA-IV genotype (R_e_ = 0.97, 95% CI: 0.72 – 1.25) and the Auckland community cluster (R_e_ = 1.09, 95% CI: 0.79 – 1.48). Infectious periods (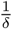, the time from acquisition to death / sampling of the strain) were estimated on the order of several years for strains from NZ (3.72 years, 95% CI: 1.12 - 6.10), NT (2.29 years, 95% CI: 0.97 – 3.41), PNG (2.21 years, 95% CI: 0.49 – 5.08) and FNQ (1.06 years, 95% CI: 0.19 - 2.58) (Table S1, Fig. S1). We note that birth-death skyline models assume well-mixed populations and that the comparatively low, lineage-wide median estimate of the infectious period in ST93 (0.427 years, 95% CI: 0.244 – 0.63, Table S1, Fig. S3) was likely a result of applying the model over the early diverging MSSA clade and the MRSA clades on the AEC, where the resulting population structure biased the parameter estimate for the infection period. As we had sufficient sample sizes for these subclades (n_NT_ = 96, n_AEC_ = 278), we applied the birth-death skyline to each clade individually, allowing us to model clade-specific changes in R_e_ over time (Fig. 2C, insets) as well as using the clade-specific method with fixed lineage-wide clock-rates (Table 1). This produced estimates for the duration of the infectious period consistent with the credible intervals of the outbreak subclades (NT MSSA 1.79 years, 95% CI: 0.77 – 3.76; AEC 1.38 years, 95% CI: 0.47 - 4.86) (Table S1). Stable circulation (R_e_ *≈* 1) on the Australian East Coast was observed following a notable spike in R_e_ shortly after acquisition of SCC*mec*-IV in the MRCA of the clade (Fig. 2C). In contrast, R_e_ of ST93-MSSA in the NT has been increasing since around 2003, with credible intervals of R_e_ > 1 suggesting sustained transmission until at least 2011. More recent genomic data on the spread of ST93-MSSA was not available.

### Sustained community transmission of the Bengal Bay clone (ST772-MRSA-V) in Pakistan

We next investigated whether clade-specific signatures of epidemic growth (R_e_ > 1) could be found in other community-associated MRSA lineages. We had previously reconstructed the detailed (n = 355) evolutionary history of the ST772-MRSA-V clone (15), which acquired multiple resistance elements, and emerged in the last two decades on the Indian subcontinent, where it has a become a dominant community-associated lineage (2). No other genomic samples were available from these countries with the exception of unreleased ST772-A samples from India (43) and a macaque-associated environmental MRSA isolate from Nepal (44). We sequenced an additional 59 strains of ST772 from community and hospital sources in the population centers of Rawalpindi and Haripur in Pakistan (39), as well as some strains imported into a University hospital in Norway (21) (Fig. 3). We found that ST772 was exported into Pakistan on multiple occasions from the background population on the Indian subcontinent (Fig. 3); our sample contained several smaller transmission clusters (n < 8) in line with observations of community spread following international transmission (15, 21) (Fig. 3A). In addition, a larger transmission cluster (n = 25) was established shortly after fixation of SCC*mec*-V (5C2) (2002, 95% CI: 2000 - 2003) in the emergent clade ST772-A2 (Fig. 3A). Application of the birth-death skyline model on the lineage revealed changes in effective reproduction numbers similar to those observed in ST93-MRSA-IV (Fig. 3B). Instead of several pronounced spikes of the reproduction number, its epidemic phase was characterized by a monotonic rise in R_e_ coinciding with the acquisition of a multidrug resistance-encoding integrated plasmid (*blaZ-aphA3-msrA-mphC-bcrAB*) around 1995 (95% CI: 1992 - 1996). Following a switch in fluoroquinolone resistance mutations in *gyrA* and fixation of the SCC*mec*-V (5C2) variant shortly after its emergence on the Indian subcontinent (1998, 95% CI: 1996 - 1999), a smaller increase in the reproduction number occurred with a delay of several years (Fig. 3B). Estimates for R_e_ in the Pakistan cluster suggest that importation resulted in sustained transmission (R_e_ = 1.36, 95% CI: 1.08 - 1.79). Mirroring the emergence of drug-resistant ST93-MRSA on the Australian East Coast, the Bengal Bay clone emerged in population centers on the Indian subcontinent from a currently unknown MSSA progenitor population and was able to establish sustained transmission after importation into Pakistan (Fig. 3C).

**Fig. 3.**
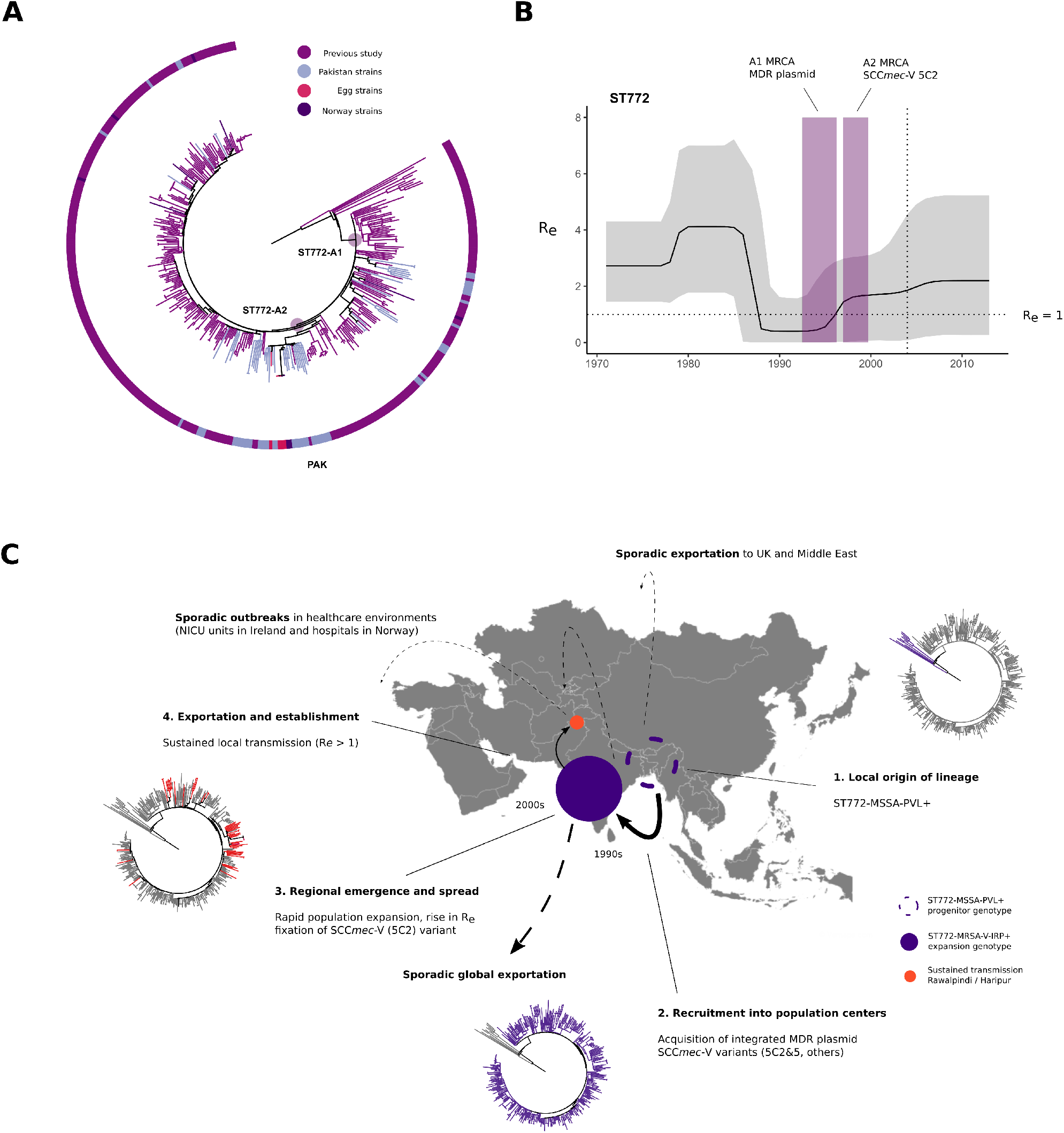
Genomic epidemiology of the Bengal Bay clone ST772 on the Indian subcontinent. (A) Rooted maximum-likelihood phylogeny of ST772 showing new strains (n = 59) from community transmission in Haripur and Rawalpindi (Islamabad metropolitan area). Sporadic importation into Pakistan is evident from singular and small transmission clusters, including a larger community transmission cluster in Rawalpindi (n = 25, PAK), where table-eggs were associated with the community outbreak and indicated additional spread overseas (B) Effective reproduction number (R_e_) over time; acquisition of the MDR integrated plasmid (MRCA 95% CI colored) is associated lineage-wide epidemic spread (R_e_ > 1). Subsequent rapid fixation of the SCC*mec*-V (5C2) within a couple of years indicates a delayed effect on R_e_ increasing slightly after the variant cassette fixation. (C) Events in the emergence of drug resistant ST772-MRSA on the Indian subcontinent; branch color in maximum likelihood phylogenies show major subclades and ongoing transmission in Pakistan. (1) Local emergence of ST772-MSSA-PVL+ in the Bengal Bay area (first samples Bangladesh and India, 2004) (2) Acquisition and chromosomal integration of a multidrug-resitant plasmid encoding *blaZ-aphA3-msrA-mphC-bcrAB* producing a ST772-MSSA-MDR clade which experiences multiple introgressions of SCCmec-V variants (5C2 and 5c2&2). This genotype successfully spreads in the wider population of the Indian subcontinent. Eventually the shorter variant SCC*mec*-V-5C2 becomes fixed in the population (3); meanwhile the pathogen population grows on the Indian subcontinent, transmission is increasing and sustained (R_e_ > 1); massive sporadic exportation from the subcontinent is occuring including into Pakistan (4), where a community outbreak establishes sustained transmission (Table 1)

### Global emergence and trajectory of community-associated *Staphylococcus aureus*

We next applied the birth-death skyline model to other community-associated MRSA clones, accounting for major lineages that have become dominant community lineages regionally and for which lineage-resolved genomic data were available (n > 100, Fig. 4) (7–15). Short-read sequence data with dates and locations from previous genomic lineage analyses were collected from studies published on the emergence of ST1 (n = 190), ST152 (n = 139) and ST80 (n = 217) in Europe, the US-Taiwan clone ST59 (n = 154) and the European-American ST8 (n = 210, excluding isolates available as assemblies only). Multiple sequence types (ST152, ST8, ST80) included extant MSSA populations circulating in Africa (Supplementary Tables, Table 1). Our analysis confirmed signatures of epidemic growth across these lineages, including notable increases in R_e_ following genomic changes and recruitment into regional host populations (Central Europe, North America, Australian East Coast, India, Taiwan), as well as increases in N_e_ (effective population sizes of *S. aureus* lineages) noted in previous investigations, coinciding with increases of R_e_ (Fig. 4, Fig. S1). MRCAs of antibiotic resistant clades in all MSSA and MRSA sublineages were estimated with 95% CI lower bounds between 1972 - 2005, and upper bounds between 1978 - 2009, confirming the seemingly convergent global emergence of resistant community strains in the late 20th century (Table 1). Low estimated sampling proportions suggest that ST8 and ST93 are widespread, consistent with global and regional epidemiological data of these clones; there was less certainty in the predictions for the recently emerged ST1, ST772 and for ST152 (Table S1). High posterior estimates of sampling proportion (*ρ*) in ST80 further suggest that the lineage is in decline in the sampled European population, although anecdotal reports indicate potential ongoing circulation in North Africa. Overall, median infectious periods varied between lineages with the shortest estimates of several months for ST93 and the longest estimates exceeding ten years in several lineages (Table S1).

**Fig. 4.**
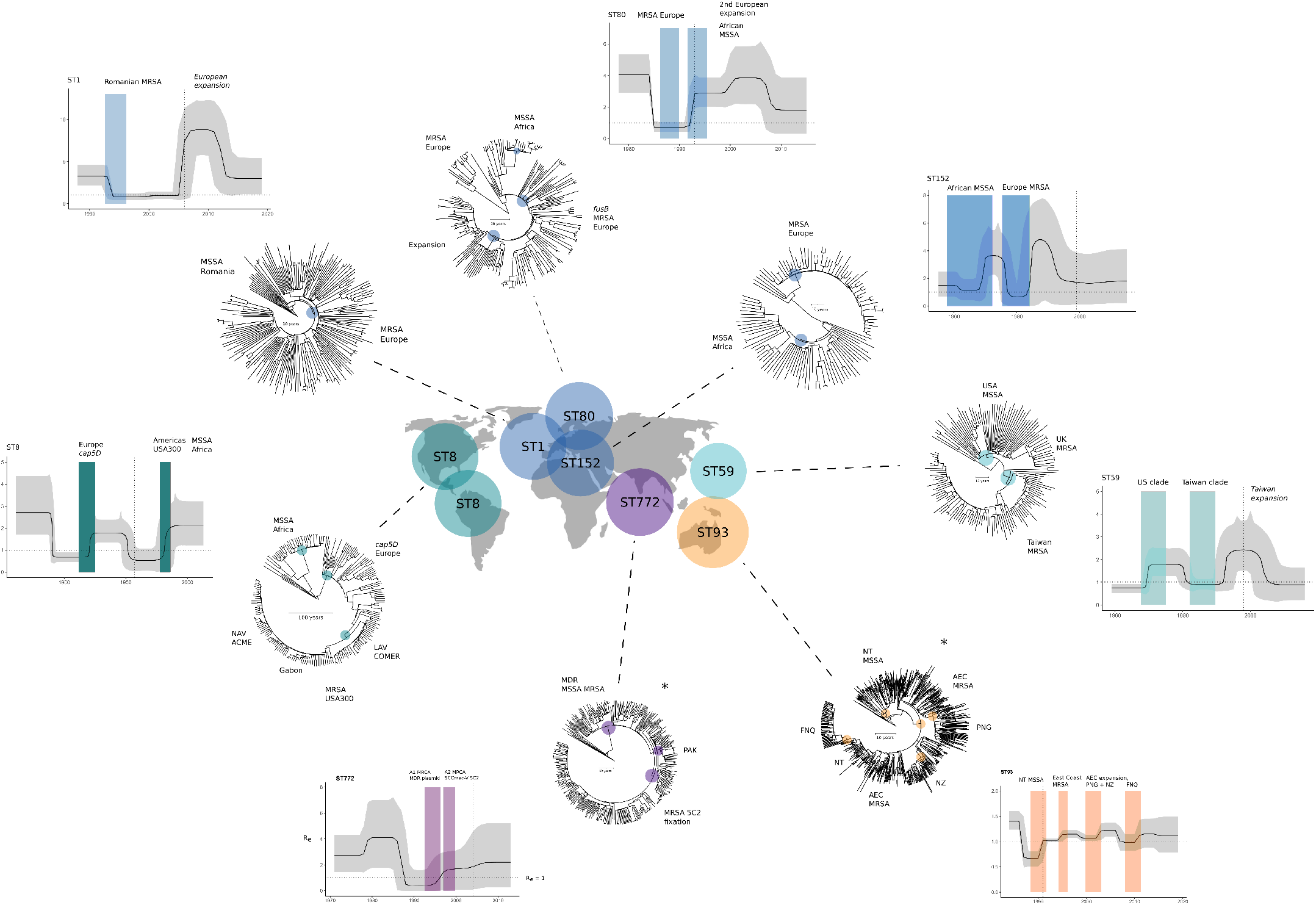
Bayesian phylogenetic trees and changes in the effective reproduction number (R_e_) of global community-associated *Staphylococcus aureus* lineages. The map shows lineages with sufficient (n > 100) genotype-resolved data included in this study, their main geographical regions in which they emerged, and their respective Bayesian maximum clade credibility trees, with colored dots indicating important clade and outbreak divergences (internal nodes). Associated changes in R_e_ estimated over equally sliced intervals over the sampling period in the birth-death skyline model are shown next to the trees. Trajectories of R_e_ show the median posterior estimates over time (dark line) and their 95% CI intervals (grey). Colored rectangles in the skyline plots correspond to 95% CI intervals of the clade MRCA indicated by dots in the lineage trees.

Considerable changes in R_e_ occurred in the ancestral ST8-MSSA genotype at the emergence in European populations in the 19th century, which has been associated with the capsule mutation *cap5D* (7) (1860, 95% CI: 1849 - 1871) (Fig. 4). The proto community-associated clone then spread to the Americas and acquired SCC*mec*-IV variants as well as the canonical COMER and ACME elements at the divergence of two regionally distinct epidemics across North America (14) and parts of South America (13, 45) which are notable as a combined increase in R_e_ in the second half of the 20th century (Fig. 4). While data was sparse for ST59-MSSA strains (9), elevated reproduction numbers indicate that it became epidemic in the United States in the 1970s and 1980s, followed by the emergence of a resistance-enriched MRSA clade in Taiwan in the late 1970s and its expansion in the 1990s with a delay between the MRCA of resistant strains and the epidemic in Taiwan several years later (Fig. 4, Table 1). Similar delays occurred in European clones ST80-MRSA and ST1-MRSA, which also shared high estimates for their infectious periods (> 10 years). We suspect that ancestral strains circulated for several years in local subpopulations before their emergence across Europe in the 1990s (ST80) (8) and 2000s (ST1) (11) but a weak temporal signal may contribute to a high degree of uncertainty in ST1 (Fig. S4, 95% CI intervals). Similar to the minor increase in R_e_ of ST772 after acquisition of the SCC*mec*-V (5C2) variant, a second increase in R_e_ without notable genomic changes was observed in ST80, suggesting a second shift in transmission dynamics as the MRSA genotype spread across Europe in the early 2000s (Fig. 4). We observed the steepest spikes of reproduction numbers in clones recruiting into European countries (ST1, ST152) where R_e_ temporarily spiked to > 5.0 - 8.0 after initial recruitment into the host population, albeit with large confidence intervals (Fig. 4). R_e_ estimates of West African subclades indicated epidemic spread in symplesiomorphic MSSA (ST8-MSSA R_e_ = 1.54, 95% CI: 1.09 - 2.26; ST152-MSSA R_e_ = 1.55, 95% CI: 1.10 – 2.25) and an introduction of USA300 (MRSA) in Gabon (ST8-MRSA-Gabon, R_e_ = 1.58, 95% CI: 1.11 - 2.34). However, these MSSA clades had acquired mild beta-lactam and other antibiotic resistance before their regional spread, including notable enrichment of *blaZ, dfrG, tetK* and *fosD* in ST8-MSSA, *blaZ* in ST152-MSSA, *blaZ* and *ermC* in ST93-MSSA, as well as occasional acquisition or loss of SCC*mec* in most MSSA clades and sublineages (Fig. S8).

## Discussion

In this study, we found a pattern in the emergence of community clones (n_total_ = 1843) associated with the acquisition of antibiotic resistance determinants, which coincide with changes in host-pathogen transmission dynamics (increases in R_e_) and lineage population size expansions (N_e_) upon recruitment into regional population centers in the 1990s. Increases in R_e_ exceeding the epidemic threshold (R_e_ > 1) were closely associated with the acquisition of resistance in community-associated MSSA circulating in specific host subpopulations. AMR acquisition was followed with the emergence and sustained transmission of resistant clades in regional population centers, such as the Australian East Coast (ST93), Taiwan (ST59), the Indian subcontinent (ST772) and central Europe (ST1, ST80, ST152). We hypothesize that resistance acquisition enables niche transitions into host populations with distinct socioeconomic structure and population densities, particularly those in urban or industrialized settings. These ‘AMR spillover’ events show patterns in R_e_ reminiscent of pathogens recruiting into susceptible host populations, where spikes in the effective reproduction numbers are followed by establishment of sustained transmission (R_e_ > 1) or elimination (R_e_ < 1). Sharp increases in R_e_ of emergent lineages were found concomitant with the MRCA of clades that had obtained AMR elements or mutations (Fig. S8). Fixation of resistance determinants and leveling of R_e_ following these spikes suggests that epidemiological factors such as widespread antibiotic use in the community, improved access to healthcare services and treatment, public health responses, or environmental antimicrobial contamination, may constitute a new adaptive landscape for the emerging drug-resisant clade, contributing to its successful dissemination or elimination. This is observed in the persistence of the epidemics (R_e_ > 1) over decades following the initial spikes in emergence, which often coincided with the first available samples of these lineages (Fig. 4, vertical lines in subplots)

Estimates of R_e_ are susceptible to a number of demographic factors which we could not explicitly model, including access to treatment, host population contact density, and changes in age-specific mixing patterns and others. Signatures in R_e_ over time inferred from these genomic data therefore combine demographic and epidemiological factors linked to genomic changes and geographical strain attribution in the jointly inferred phylogenies. Our data suggest that the acquisition of multiple locality-specific resistance mutations and mobile genetic elements has driven rapid genotype expansions with notable increases in R_e_ observed across lineage-wide and clade-specific analyses. SCC*mec*-elements of type IV and V eventually integrated into resistant clades, but cassette genotypes are variable and usually preceded or supplemented by other resistance determinants. For example, the stepwise acquisition of resistance in ST772-MSSA occured first through the chromosomal integration of a multidrug-resistance plasmid, followed by a shift in the *gyrA* mutation conferring fluoroquinolone resistance and eventual fixation of the short 5C2 variant of SCC*mec*-V (15), whereas other lineages such as ST93-MRSA-IV emerged after a singular acquistion event of SCC*mec*-IV (10). It is notable that even the successful and sampled MSSA clades in Africa and northern Australia were enriched in resistance determinants, with *blaZ* (ST93-MSSA, ST8-MSSA, ST152-MSSA) and *tetK* (ST8-MSSA) amongst others (Fig. S8).

In support of the “AMR spillover” hypothesis, Gustave and colleagues (46) demonstrated in competition experiments with ST8 (USA300) and ST80 genotypes, that antibiotic-resistant strains expressed a fitness advantage over wild-type strains on subinhibitory antibiotic media. Presence of low-level antibiotic pressure may therefore be a crucial epidemiological driver in the emergence of resistant clades in host populations that have widespread access to treatment or may not practice effective antibiotic stewardship. However, antibiotic resistance is likely not the only driver for local clade emergence and dissemination. Gustave and colleagues (45) recently showed that the mercury-resistance operon located on the COMER element may have driven the dissemination of the USA300 variant in South America on a background of pollution from mining activities. Further data to investigate local competitive fitness dynamics under population antibiotic pressure backgrounds *in vivo* or *in vitro* is required and has been laid out in experimental work (45). Complex genotype competition dynamics may arise from environmental coupling at different time-points in the evolution of a lineage, and differential competitive fitness in evolutionary and epidemiological landscapes may play a role in why some community strains successfully recruit into host populations following exportation and fail to become endemic elsewhere.

Local epidemiological patterns in the ST93 lineage phylogeny revealed co-circulating MSSA and MRSA genotypes in the Northern Territory, where a potentially sustained (R_e_ = 0.97, 95% CI: 0.72 - 1.25) re-introduction of the MRSA genotype eventually spread into communities across Far North Queensland. We further observed that sustained transmission is occurring in symplesiomorphic MSSA populations of ST8 and ST152 in Africa (7, 12), as well as extant ST93-MSSA in northern Australia, particularly amongst Indigenous communities (10, 28, 47). MSSA clades thus have established sustained transmission in host populations, preceding MRSA clade recruitment into geographically distinct populations (ST8-USA300, ST93-MRSA-IV, ST152-MRSA). It is notable that epidemic signatures were found for resistant MSSA (Northern Territory, Africa) and MRSA (FNQ, PNG, NZ) clades in socioeconomic settings similar to those experienced by many remote Indigenous communities in Australia. These include high burdens of skin-disease, domestic overcrowding, and poor access to healthcare or other public services (10, 47– 50).

Non-synonymous mutations in factors associated with immune response and skin colonization at the divergence of epidemic MRSA and MSSA have been detected previously in ST772 (15), and ACME and COMER elements in the USA300 clades are implicated in transmission and persistance phe-notypes (13, 51, 52), but it is unclear to what degree these changes have contributed to the emergence, transmission, persistence, or fitness of resistant strains in the presence of other strains or genotypes. Given that acquisition of antibiotic resistance determinants and recruitment into population centers coincides with rapid increases in R_e_ across all lineages examined in this study, these factors are not likely to explain the rapid change in transmission dynamics we estimated at the divergence of resistant clades (Fig. S8). However, mutations may contribute to ongoing persistence in host populations before and after emergence, or constitute pre-adaptations that support successful transmission in new host populations. Canonical mutations have previously been detected at the divergence of ST772-A and have been associated with colonization ability, which may play a role in compensating for the fitness cost induced by resistance acquisition (15). Preliminary phenotypic data from the Bengal Bay clone suggests that there was no significant difference in biofilm formation or growth rate between MSSA and multidrug-resistant MRSA strains (15). Further rigorous experiments will need to be conducted to better understand the significance of these mutations in strains preceding the emergence of resistant clades and their interaction with resistance phenotypes.

Our sampling design and models used for the inference of phylodynamic parameters have important limitations. First, we note that uncertainty deriving from incomplete lineage sampling is large, but mitigated by using published collections of metadata-complete and lineage-representative genomes. However, there was a lack of data on ancestral MSSA strains, a problem pointed out explicitly for ST80 (8) and ST59 (9), but also relevant to ST772 (15). These effects appeared less severe for ST8, for which there was a wide sampling range going back to 1953 (7), and for ST93 (10, 16), which had well-represented MSSA collections from the Northern Territory, and for which the MRCA and origin of the lineage were estimated to have occurred within a year (Fig. S3). For our phylodynamic comparison we addressed sampling bias towards the present by allowing piecewise changes in the sampling proportion consistent with sampling effort for each lineage.

Our study provides phylogenetic and population genomic evidence that community-associated genotypes have emerged in regional host populations following the acquisition of antibiotic resistance. Pre-adaptations for transmission in the ancestral host populations may have contributed to the eventual, epidemic spread of resistant strains in populations with access to antibiotic treatment and healthcare services. However, well-sampled ancestral MSSA genomes are lacking for important community-associated lineages including ST772, ST59 and ST80; deeper sampling and ongoing genome-informed surveillance of these populations will be required to further understand the processes that allow lineages to emerge and become epidemic. It is notable that the seemingly convergent emergence events in the second half of the 20th century — whose signatures we detect from phylodynamic models across all sequence types — were caused by epidemics of both, drug-resistant MSSA and MRSA. While antibiotic resistance has facilitated emergence in regional population centers, local evolutionary and host population dynamics have also played a role in the emergence of important community-associated clades, including the potential role of environmental pollution and the COMER element in the dissemination of USA300 in South America (45).

Despite the limitations of this study, the phylodynamic estimates observed across sequence types are remarkably consistent with decades of molecular and epidemiological work that has characterised the global emergence and spread of community-associated *S. aureus* lineages, including their notable emergence in the 1990s and subsequent establishment across their respective geographical distributions. We provide genomic evidence for sustained transmission of MRSA strains in Pakistan and PNG, as well as for drug-resistant MSSA strains in African countries and northern Australia, where - in addition to the notable enrichment of resistance without stable acquisition of SCC*mec* - sociodemographic host-factors may play an under-appreciated role in the transmission dynamics and epidemic potential of these lineages. Ongoing epidemic transmission of ST93-MSSA is of concern for Indigenous communities in the Northern Territory and there is now evidence for the dissemination of its emergent MRSA genotype beyond the Australian continent. Wider circulation of ST93-MRSA-IV in Papua New Guinea is likely. Our work underlines the importance of considering remote and disadvantaged populations in a domestic and international context. Social and public health inequalities (28, 53) appear to facilitate the emergence and circulation of community-associated pathogens including drug-resistant *S. aureus*.

## Materials and Methods

### Outbreak sampling and sequencing

We collected isolates from outbreaks in two remote populations in northern Australia and Papua New Guinea (Fig. 1). Isolates associated with paediatric osteomyelitis cases (mean age of 8 years) were collected from 2012 to 2017 (n = 42) from Kundiawa, Simbu Province (27), and from 2012 to 2018 (n = 35) from patients in the neighbouring Eastern Highlands province town of Goroka. We supplemented the data with MSSA isolates associated with severe hospital-associated infections and blood cultures in Madang (Madang Province) (n = 8) and Goroka (n = 12). Isolates from communities in Far North Queensland, including metropolitan Cairns, the Cape York Peninsula and the Torres Strait Islands (n = 91), were a contemporary sample from 2019. Isolates were recovered on LB agar from clinical specimens using routine microbiological techniques at Queensland Health and the Papua New Guinea Institute of Medical Research (PNGIMR). Isolates were transported on swabs from monocultures to the Australian Institute of Tropical Health and Medicine (AITHM Townsville) where they were cultured in 10 ml LB broth at 37°C overnight and stored at -80°C in glycosol and LB. Samples were regrown prior to sequencing, and a single colony was placed into in-house lysis buffer and sequenced at the Doherty Applied Microbial Genomics laboratory using 100 bp paired-end libraries on Illumina HiSeq. Illumina short-read reads from the global lineages included in this study were collected from the European Nucleotide Archive (Supplementary Tables).

### Genome assembly and variant calling

Illumina data was adapter- and quality-trimmed with `Fastp` (54) before *de novo* assembly with `Shovill` (https://github.com/tseemann/shovill) using downsampling to 100x genome coverage and the Skesa assembler (55). Assemblies were genotyped with `SCCion` (https://github.com/esteinig/sccion), a wrapper for common tools used in *S. aureus* genotyping from reads or assemblies. These include multilocus sequence (MLST), resistance and virulence factor typing with `mlst` and `abricate` (https://github.com/tseemann) using the `ResFinder` and `VFDB` databases (56, 57). SCC*mec* types were called using the best Mash (58) match of the assembled genome against a sketch of the `SCCmecFinder` database (59) and confirmed with *mecA* gene typing from the `ResFinder` database. Antibiotic resistance to twelve common antibiotics were typed with `Mykrobe` (53); this strategy was used for all lineage genomes to confirm or supplement (7) antibiotic resistance determinants as presented in original publications. Strains belonging to ST93 (FNQ, PNG) and ST772 (Pakistan) were extracted and combined with available sequence data from previous studies on the ST772 (n_total_ = 359) and the ST93 (n_total_ = 575) lineages. `Snippy` v.4.6.0 (https://github.com/tseemann/snippy) was used to call core-genome SNPs against the ST93 (6,648 SNPs) or ST772 (7,246 SNPs) reference genomes JKD6159 (60) and DAR4145 (20). Alignments were purged of recombination with `Gubbins` (61) using a maximum of five iterations and the *GT R* + *G* model in (40). Quality control, assembly, genotyping, variant calling and maximum likelihood (ML) tree construction, statistical phylodynamic reconstruction and exploratory Bayesian analyses were implemented in Nextflow (62) for reproducibility of the workflows (https://github.com/np-core/phybeast). Phylogenetic trees and metadata were visualized with Interactive Tree of Life (63). All program versions are fixed in the container images used for analysis in this manuscript (Data Availability).

### Maximum-likelihood phylogenetics and -dynamics

We used a ML approach with `TreeTime` v0.7.1 (41) to obtain a time-scaled phylogenetic tree by fitting a strict molecular clock to the data (using sampling dates in years throughout). Accuracy for an equivalent statistical approach using least-squares dating (64) (LSD) is similar to that obtained using more sophisticated Bayesian approaches with the advantage of being computationally less demanding (65). As input, we used the phylogenetic tree inferred using ML in `RAxML-NG` after removing recombination with `Gubbins`. The molecular clock was calibrated using the year of sample collection (i.e. heterochronous data) with least-squares optimization to find the root, while accounting for shared ancestry (covariation) and obtaining uncertainty around node ages and evolutionary rates. We also estimated the ML piecewise (skyline) coalescent on the tree using default settings, which provides a baseline estimate of the change in effective population size (N_e_) over time. Temporal structure of the data was assessed by conducting a regression of the root-to-tip distances of the ML tree as a function of sampling time and a date-randomisation test on the TreeTime estimates with 100 replicates (65, 66) (Fig. S1). All trees were visualized in Interactive Tree of Life (ITOL) and node-specific divergence dates extracted in Icytree.

### Bayesian phylodynamics and prior configurations

We used the Bayesian coalescent skyline model to estimate changes in effective population size (N_e_), and implemented the birth-death skyline from the `bdsky` package (https://github.com/laduplessis/bdskytools) for `BEAST` v2.6 to estimate changes in the effective reproduction number (R_e_) (33, 34, 42). Birth-death models consider dynamics of a population forward in time using the (transmission) rate *λ*, the death (become uninfectious) rate *δ*, the sampling probability *ρ*, and the time of the start of the population (outbreak; also called origin time) *T*. The effective reproduction number (R_e_), can be directly extracted from these parameters by dividing the birth rate by the death rate (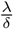).

For lineage-wide analysis we used all available samples from each sequence type, including MSSA and MRSA clades, but compared model estimates from the entire lineage to distinct clade subsets to mitigate the effect of population structure from well-sampled clones like ST93 (Table 1, Fig. 2B: R_e_ estimates for ST93-MSSA and -MRSA). Median parameter estimates with 95% CI intervals and Markov chain traces were inspected in `Tracer` to assess convergence (67). We implemented Python utility functions to generate XML files and configure priors more conveniently in a standardized form (`nanopath beastling`) implemented in the `NanoPath` package (https://github.com/esteinig/nanopath). Plots were constructed with scripts (https://github.com/esteinig/publications) that use the bdkytools package including computation of the 95% highest posterior density (HPD) median intervals (credible intervals, CI) using the Chen and Shao algorithm implemented in the boa package for R (68). We chose to present median posterior intervals, since some posterior distributions had long tails in the prior distributions of some parameters (e.g. origin or become uninfectious rates, Fig. S3). Icytree (69) was used to inspect Bayesian maximum clade credibility trees derived from the posterior sample of trees. In the coalescent skyline model for each lineage, we ran exploratory chains of 200 million iterations varying the number of equidistant intervals over the tree height (dimensions, *d*) specified for the priors describing the population and estimated interval (dimension) size (*d* = *{*2, 4, 8, 16 *}*, Fig. S2). As posterior distributions were largely congruent (top rows, Fig. S2), we selected a sufficient number of intervals to model changes in effective population size (N_e_) of each lineage over time (*d* = 4 and *d* = 8).

In the birth-death skyline models, priors across lineages were configured as follows: we used a *Gamma*(2.0, 40.0) prior for the time of origin parameter (*T*), covering the last hundred years and longer. We chose a *Gamma*(2.0, 2.0) prior for the reproductive number, covering a range of possible values observed for *S. aureus* sequence types in different settings (70–73) which may have occured over the course of lineage evolution. We configured the reproduction number prior (R_e_) to a number of equally sized intervals over the tree; a suitable interval number was selected by running exploratory models for each lineage with 100 million iterations from *d* = 5*-*10 followed by a comparison of parameters estimates under these configurations (occurence of stable posterior distributions, absence of bi or multi-modal posteriors) (not shown, available in data repository). Because sequence type-specific becoming-non-infectious rates in community-associated *S. aureus* are not well known, either from long-term carriage studies or phylogenetic reconstructions (22, 74–76), we explored a range of prior configurations for the becoming uninfectious rate parameter (*δ*) including a flat uniform prior *U nif orm*(1.0, 1.0) and a *Lognormal*(*µ*, 1.0) prior with *µ* = 0.1 (10 years infectious period), *µ* = 0.2 (5 years) and *µ* = 1.0 (1 year). We chose a *Lognormal*(1.0, 1.0) prior, as the resulting parameters estimates were coherent (Fig. S5). Lineage-wide sensitivity analysis showed that estimates across lineages were not driven by the prior (Fig. S6). Sampling proportion (*ρ*) was fixed to zero in the interval ranging from the origin to the first sample (pre-sample period); the remaining time until present (sampling period) was estimated under a flat *Beta*(1.0, 1.0) prior, accounting for sampling bias towards the present as well as largely unknown estimates of global sampling proportions across lineages. Final lineage models were run with 500 million iterations on GPUs with the `BEAGLE` library (77) under a *GT R* + *Gamma* substitution model with four rate categories. We used a strict molecular clock with a *Lognormal*(0.0003, 0.3) rate prior in real space as all lineages ‘evolved measurably’ (Fig. S1). Models were run until chains were mixed and ESS values reached at least 200, as confirmed in `Tracer`.

Lastly, we ran birth-death skyline models on specific clades within the lineage phylogenies, including the ancestral and symplesiomorphic MSSA populations, the USA300 sublineages, and importations of ST93, ST772 and ST8 (Fig. 4, Fig. S8). For each subset of strains, we extracted the core-genome variant alignment subset, configured the reproduction number prior to a single estimate over the clade (since in outbreak datasets the number of sequences per clade was smaller than 100 and the sampling interval smaller than 10 years) (Fig. S8). Because temporal signal is lost in the clade subsets, we fixed the substitution rate to the lineage-wide estimate in all runs for 100 million iterations for the MCMC with trees sampled every 1000 steps. Runs were quality controlled by assuring that chains mixed and ESS values for all posterior estimates reached at least 200. Since sufficient samples and a wide sampling interval were available to track R_e_ changes in ST93-MSSA (n = 116 in the Northern Territory) and -MRSA clades (n = 278, Australian East Coast) over time (Fig. 2, inset plots), we explored a stable configuration of the reproducuctive number prior across equally-spaces intervals (*d* = 5 *-*10) with 200 million iterations of the MCMC (not shown, available in data repository). We explored *Gamma*(2.0, *θ*) distributions where *θ ∈ {* 0.5, 1.0, 1.5, 2.0*}* in the R_e_ prior of sublineages and outbreaks. This was to guard against bias towards inferring sustained transmission (R_e_ > 1) in outbreak models with limited data and temporal signal from subset alignments (representative examples: Fig. S7). Results from the model runs under the conservative *Gamma*(2.0, 0.5) prior assert only minor differences compared to higher configurations (consistently R_e_ > and the conservative estimates are presented here (Table 1). We also conducted a sensitivity analysis for all sublineage models where we ran the models under the prior only (Fig. S8, note that under-the-prior posteriors are still influenced by dates, albeit not the genetic data). With the excception of ST93-NT and ST93-NZ we asserted that estimates of R_e_ were not driven by the prior and dates alone in all sublineages and outbreaks. Full explorative data can be found in the data repository for this study.

## Data Availability

Sequencing data can be found at BioProject PRJNA657380. XML model files, log files of `BEAST2` runs, and links to command line interfaces for replicating the analyses and plots in this manuscript can be found at: https://github.com/esteinig/ca-mrsa

## ACKNOWLEDGMENTS

ES is supported by a PRISM2 & HOT North pilot grant (78) (NHMRC ID: 1131932). CF is supported by a HOT North fellowship (NHMRC ID: 1131932). SYCT is supported by an Australian NHMRC fellowship (ID 1145033). GPU models were run on the LIEF HPC-GPGPU Facility hosted at the University of Melbourne (LIEF Grant ID: LE170100200) (79).

## Supplementary Discussion

### Distribution of ST93 in Papua New Guinea

We used whole-genome sequencing data to reconstruct the evolutionary history of the first *S. aureus* genomes from Papua New Guinea, and discovered that a paediatric osteomyelitis outbreak in the remote Highlands Provinces (31) was caused by the Australian clone ST93-MRSA-IV. Multiple lines of evidence support a wider distribution of ST93 in PNG, including two discernible introductions, sustained and long-term transmission of the outbreak since the early 2000s, occurrence of two MLST allele variants, and a heterogeneous pattern of dissemination in the remote highland towns (Figs. 1, 2). It remains unclear to what extent ST93-MRSA-IV has disseminated in PNG. Our data further show that ST93-MRSA-IV is widespread in Far North Queensland and is likely the cause for the increasing rates of MRSA observed in FNQ communities over the last decade (29, 30). It is unclear what is driving the local persistence and evolution of the ST93-MRSA-IV genotype in PNG, and reservoirs in the community remain to be investigated. Data on antibiotic consumption in Simbu Province or Eastern Highland Province was not available. While antibiotic stewardship may play a role in the dissemination of ST93 in FNQ (29), and ST772 in the Islamabad-Rawalpindi metropolitan area (39), sustained circulation of virulent and transmissible clones in remote settings like PNG may also have been a result of historical transmission opportunities from the Australian East Coast after the emergence of ST93-MRSA-IV, as well as existing strain diversity and competitive interactions in the highlands, even in the absence of widespread antimicrobial consumption.

### Model parameters and local transmission dynamics

Our lineage-wide phylodynamic census provides genomic evidence of successful recruitment of ST93-MRSA-IV into domestic and international host populations. Estimates from Bayesian phylodynamic models indicate that sustained transmission (R_e_ > 1) following importation has not only occurred in PNG and FNQ (ST93) but also in Pakistan (ST77-MRSA-V), several African countries (ST152-MSSA, ST8-USA300), South America (ST8-USA300 COMER variant) and Europe (ST8-USA300). It therefore appears that community-associated MRSA lineages are able to establish sustained transmission after dissemination of resistant genotypes.

**Table S1.** Birth-death skyline median posterior estimates (cont’d from Table 1)

| Lineage | $T$ | 95% CI | $\frac{1}{\delta}$ | 95% CI | $\rho$ | 95% CI |
| --- | --- | --- | --- | --- | --- | --- |
| **Sequence types** |  |  |  |  |  |  |
| ST93 | 1982 | 1982 - 1983 | 0.47 | 0.24 - 0.63 | 0.003 | 0.0008 - 0.006 |
| ST772 | 1967 | 1962 - 1972 | 7.81 | 1.89 - 15.02 | 0.22 | 0.013 - 0.56 |
| ST80 | 1976 | 1974 - 1978 | 12.07 | 8.57 - 15.51 | 0.849 | 0.56 - 0.99 |
| ST8 | 1855 | 1844 - 1865 | 10.61 | 4.05 - 21.33 | 0.308 | 0.004 - 0.86 |
| ST1 | 1986 | 1984 - 1988 | 12.93 | 7.21 - 19.14 | 0.49 | 0.16 - 0.94 |
| ST59 | 1955 | 1949 - 1958 | 2.93 | 0.95 - 5.72 | 0.005 | 0.0002 - 0.018 |
| ST152 | 1951 | 1944 - 1957 | 12.69 | 5.37 - 22.71 | 0.14 | 0.01 - 0.52 |
| **MRSA sublineages** |  |  |  |  |  |  |
| ST93-MRSA-EastCoast | 1991 | 1981 - 1995 | 2.72 | 0.504 - 6.27 | 0.11 | 0.002 - 0.407 |
| ST93-MRSA-FNQ | 2005 | 2000 - 2008 | 1.06 | 0.19 - 2.58 | 0.03 | 0.004 - 0.096 |
| ST93-MRSA-PNG | 1998 | 1990 - 2002 | 2.21 | 0.49 - 5.08 | 0.01 | 0.0001 - 0.045 |
| ST93-MRSA-NT | 1998 | 1994 - 2001 | 2.29 | 0.97 - 3.40 | 0.61 | 0.15 - 0.99 |
| ST93-MRSA-NZ | 2000 | 1994 - 2000 | 3.72 | 1.13 - 6.11 | 0.51 | 0.09 - 0.99 |
| ST772-MRSA-Pakistan | 1999 | 1989 - 2003 | 1.25 | 0.41 - 2.58 | 0.03 | 0.002 - 0.15 |
| ST152-MRSA-Europe | 1985 | 1974 - 1991 | 4.07 | 0.65 - 10.34 | 0.05 | 0.0003 - 0.296 |
| ST8-MRSA-NorthAmerica | 1980 | 1972 - 1986 | 2.26 | 0.48 - 5.11 | 0.003 | 0.000002 - 0.03 |
| ST8-MRSA-SouthAmerica | 1975 | 1964 - 1982 | 2.75 | 0.49 - 6.75 | 0.001 | 0.000001 - 0.01 |
| ST8-MRSA-France |  |  |  |  |  |  |
| ST8-MRSA-Gabon | 1990 | 1981 - 1995 | 2.52 | 0.41 - 6.16 | 0.009 | 0.00001 - 0.07 |
| ST59-MRSA-Taiwan | 1974 | 1961 - 1980 | 4.25 | 0.64 - 10.45 | 0.03 | 0.0004 - 0.16 |
| **MSSA sublineages** |  |  |  |  |  |  |
| ST93-MSSA-Australia | 1986 | 1974 - 1991 | 3.01 | 0.57 - 7.17 | 0.09 | 0.0009 - 0.385 |
| ST8-MSSA-WestAfrica | 1975 | 1964 - 1982 | 3.09 | 0.52 - 7.49 | 0.0047 | 0.000002 - 0.03 |
| ST152-MSSA-WestAfrica | 1972 | 1960 - 1977 | 3.12 | 0.53 - 7.22 | 0.001 | 0.000001 - 0.009 |

Phylodynamic models predicted lineage- and clade-specific infectious periods (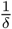) that suggest prolonged durations of infection over several years in concordance with long-term cohort studies with lineage-resolved data (Table S1). Variation in our model estimates could reflect differential local modes of persistence in the host or community (70, 80), and is susceptible to factors that we were not able to explicitly model, including access to healthcare services and treatment amongst others. It should be noted that the lineage-wide averages of infectious periods may not reflect the considerable heterogeneity in carriage duration that likely arises from the distribution of permanent and transient carriers across the population (75, 76). Our data further suggest that changes in transmission dynamics (R_e_) can occur without additional genomic changes either after incursion into a new population (e.g. ST80 European expansion; introductions of ST93 clades into PNG, FNQ and NZ) or following a delay of several years after introduction (e.g. ST772 after SCC*mec*-V (5C2) fixation, ST59 expansion in Taiwan). It is feasible that delayed changes in R_e_ may be indicative of local competitive interactions with prevailing lineages or changes in human population dynamics (e.g. in healthcare or social policies, travel and immgration policies, opening of markets and borders) that drive further dissemination once established in the host population. In one case, a sharp increase in R_e_ occurred nearly a decade after the MRCA of the resistant European ST1-MRSA clade, with the implication that the emerging genotype circulated undetected in South-East Europe (likely in Romania where the first samples originate (11, 17, 18)) before its emergence across Europe (Fig. 4).

### Additional model limitations and runtime improvements

Birth-death skyline models assume that populations are well-mixed, but clear population structure is evident between the MSSA and MRSA strains as well as in other monophyletic clades, such as the introductions of ST93 into PNG, FNQ and NZ. While we attempted to employ the more parameter-rich multitype birth-death modcels that are inherently capable of accounting for structured populations (81), the MCMC chains ultimately did not converge. This was likely due to a combination of large bacterial genome data sets and the parameter-rich model. We therefore employed the birth-death skyline model on monophyletic subsets of the lineage-wide variant alignment, provided sufficient isolates were available (Figs. 2B, 3B, Supplementary Fig. 2), thus reducing the potential impact on lineage-wide estimates arising from excessive population structure in the tree. We further explored a range of realistic configurations on the becoming uninfectious rate prior (1 - 10 years infectious period), for which few data are available from long-term surveillance studies, as well as weighting the reprodutive number prior distribution at different levels of transmission (Methods). Prior sensitivity analysis for each lineage and clade confirms that estimates were largely driven by the available data, rather than by the prior configurations (Figs. S7, S8. Further improvements to enable MCMC convergence for phylodynamic models used in large bacterial populations, including multitype birth-death models supporting larger sample sizes with numeric stability (82) and Metropolis-coupled MCMC chains (83) will be required for further investigations into the origins of community-associated *S. aureus*.

**Fig. S1.**
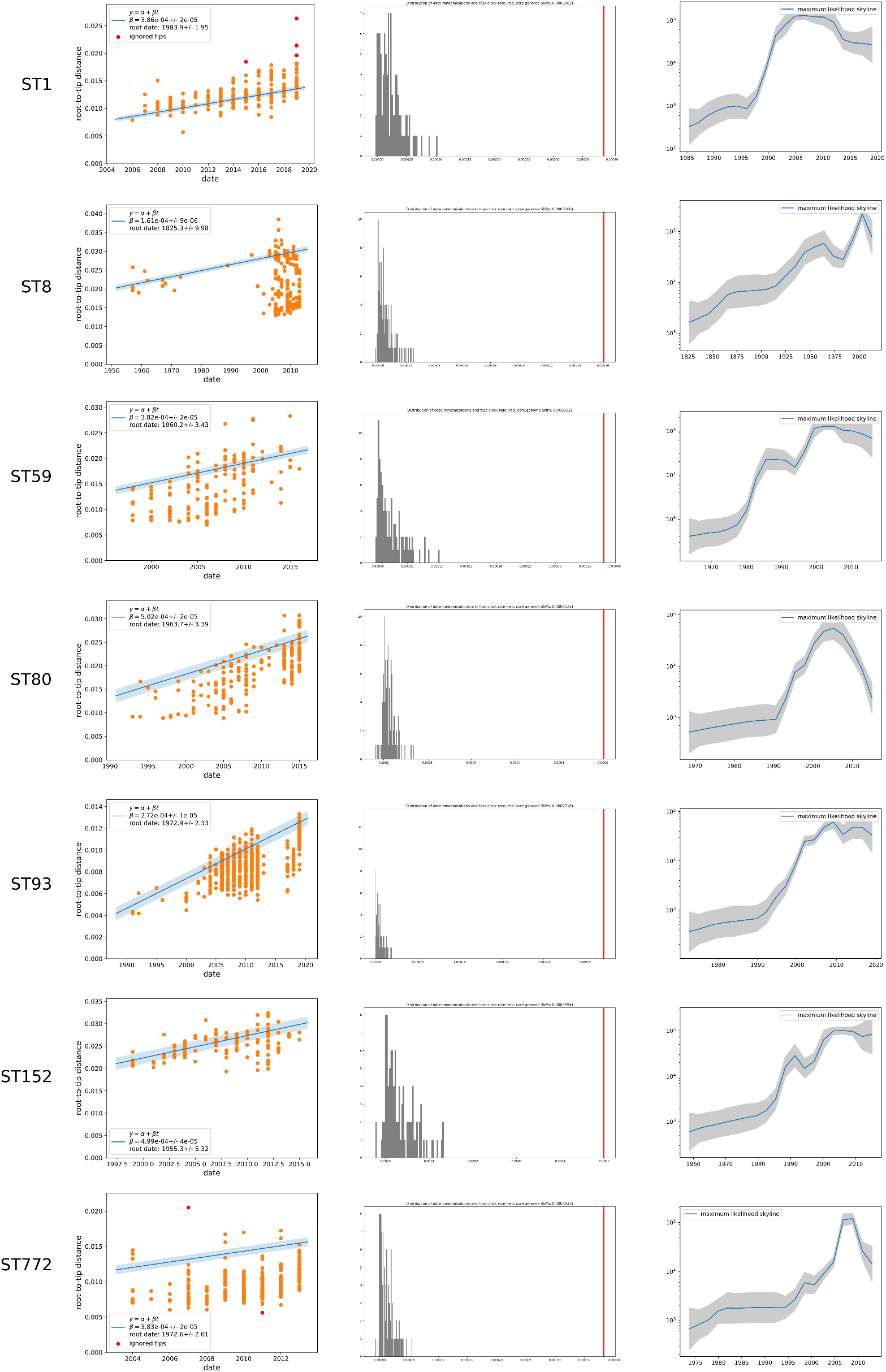
Lineage specific maximum-likelihood phylodynamics using `Treetime` (41): on the left is the root-to-tip regression on sampling dates showing that all lineages could be considered ‘measurably evolving’; in the middle the results of a date-randomisation test with 100 replicates (65, 66) indicating that all estimated clock rates (red line) are distinct from a distribution of rates estimated after randomising dates across the phylogenetic tree (gray distribution); on the right is the coalescent skyline estimate of changes in the effective population size (N_e_) over time.

**Fig. S2.**
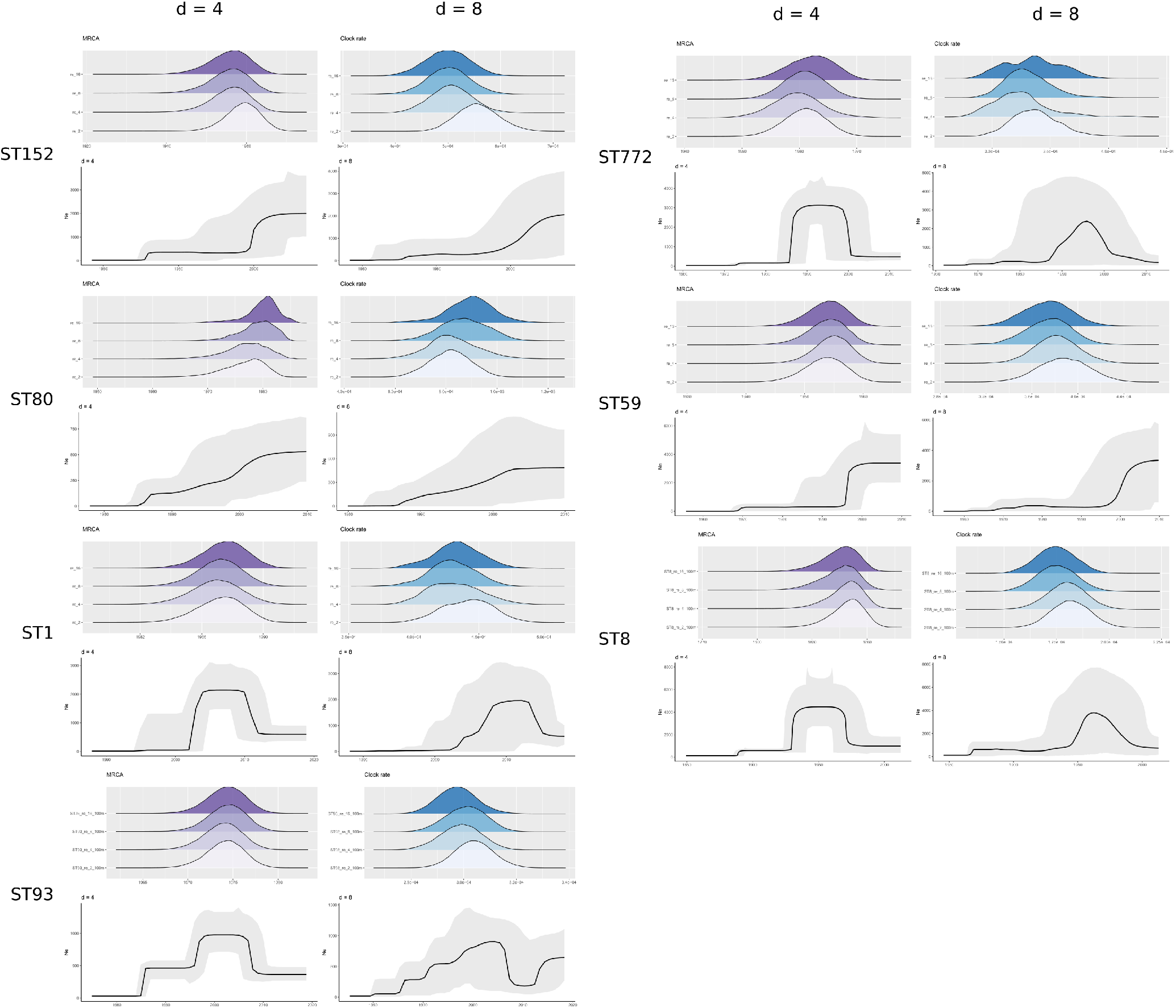
Coalescent Bayesian skyline plots of changes in the effective population size (N_e_) of *Staphylococcus aureus* lineages at different dimensional configurations (*d* = 4 and *d* = 8, skyline plots) with posterior distributions of dimensional configurations *d ∈ {*2, 4, 8, 16*}* of MRCA (purple) and clock rate (blue) above for reference.

**Fig. S3.**
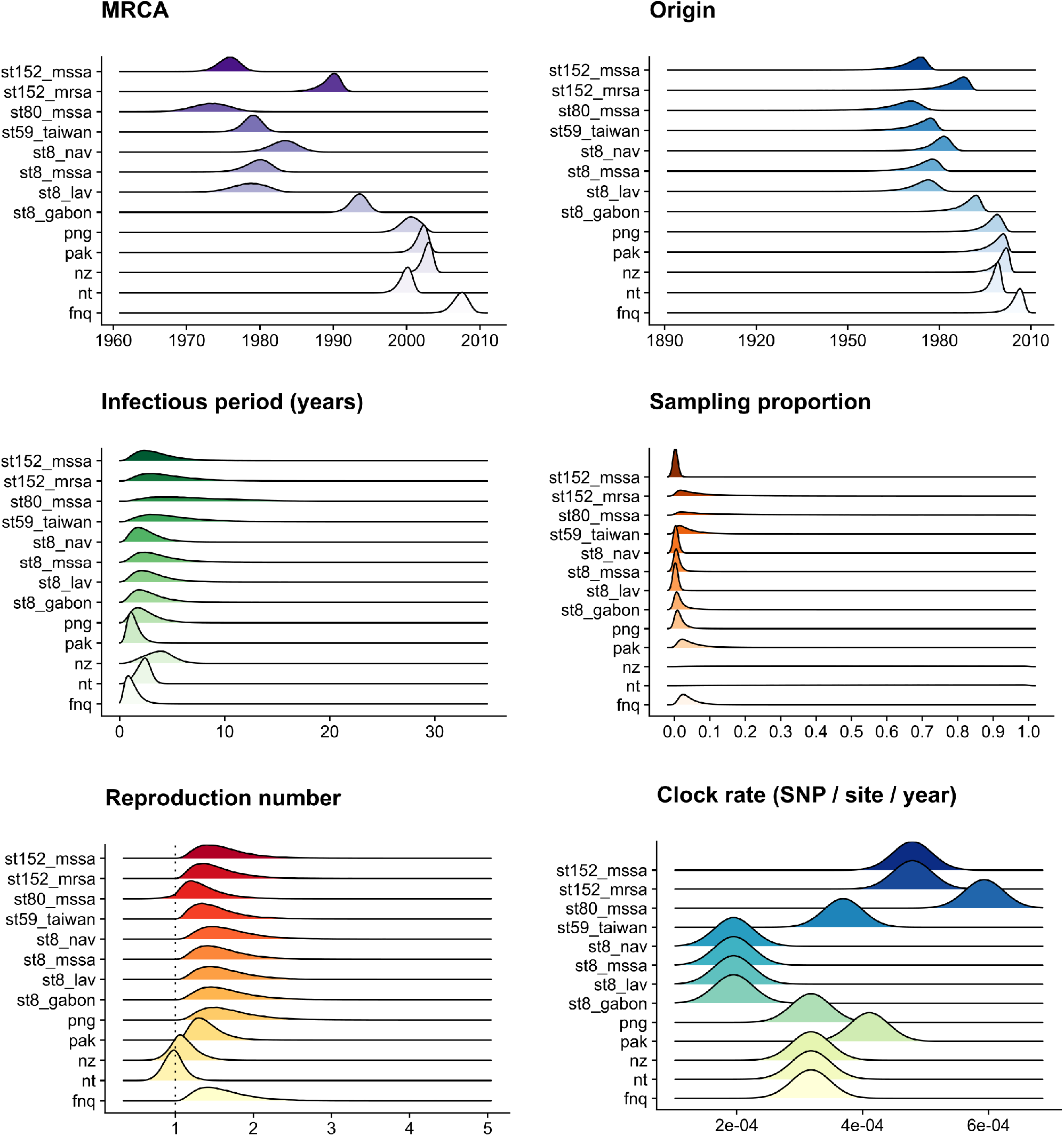
Birth-death skyline posteriors of outbreaks and sublineages in community-associated *Staphylococcus aureus*. Clock rates are fixed to the lineage-wide median higher posterior density interval (Table 1).

**Fig. S4.**
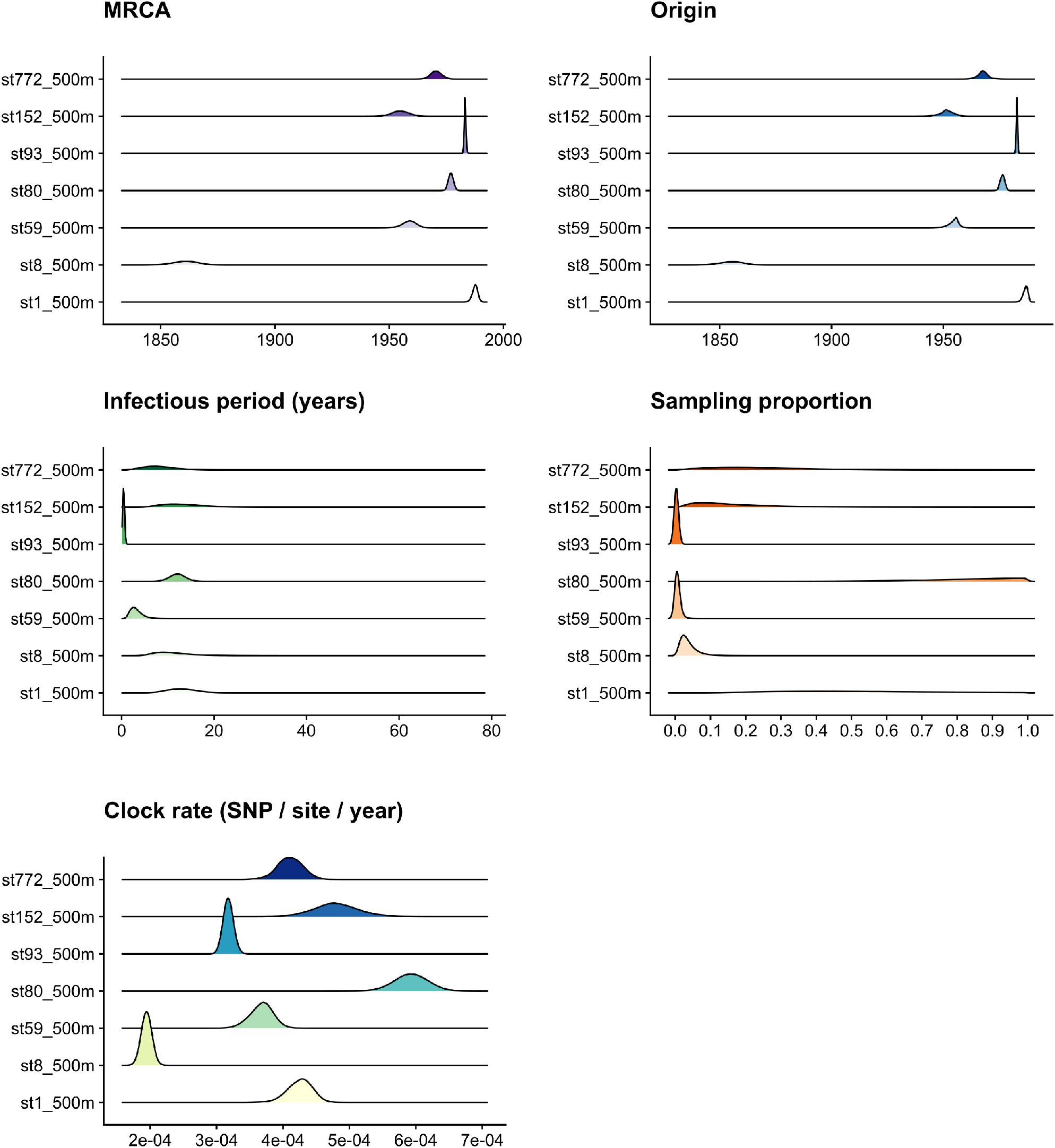
Birth-death skyline posteriors of community-associated *Staphylococcus aureus* lineages; clock rates are estimated, the sampling proportion (*δ*) prior is sliced into pre-sampling and sampling intervals, and the reproduction number (*R*e) prior varies across equally distant slices over the phylogeny (Fig. 44, Table 1, models were run for 500 million iterations of the MCMC)

**Fig. S5.**
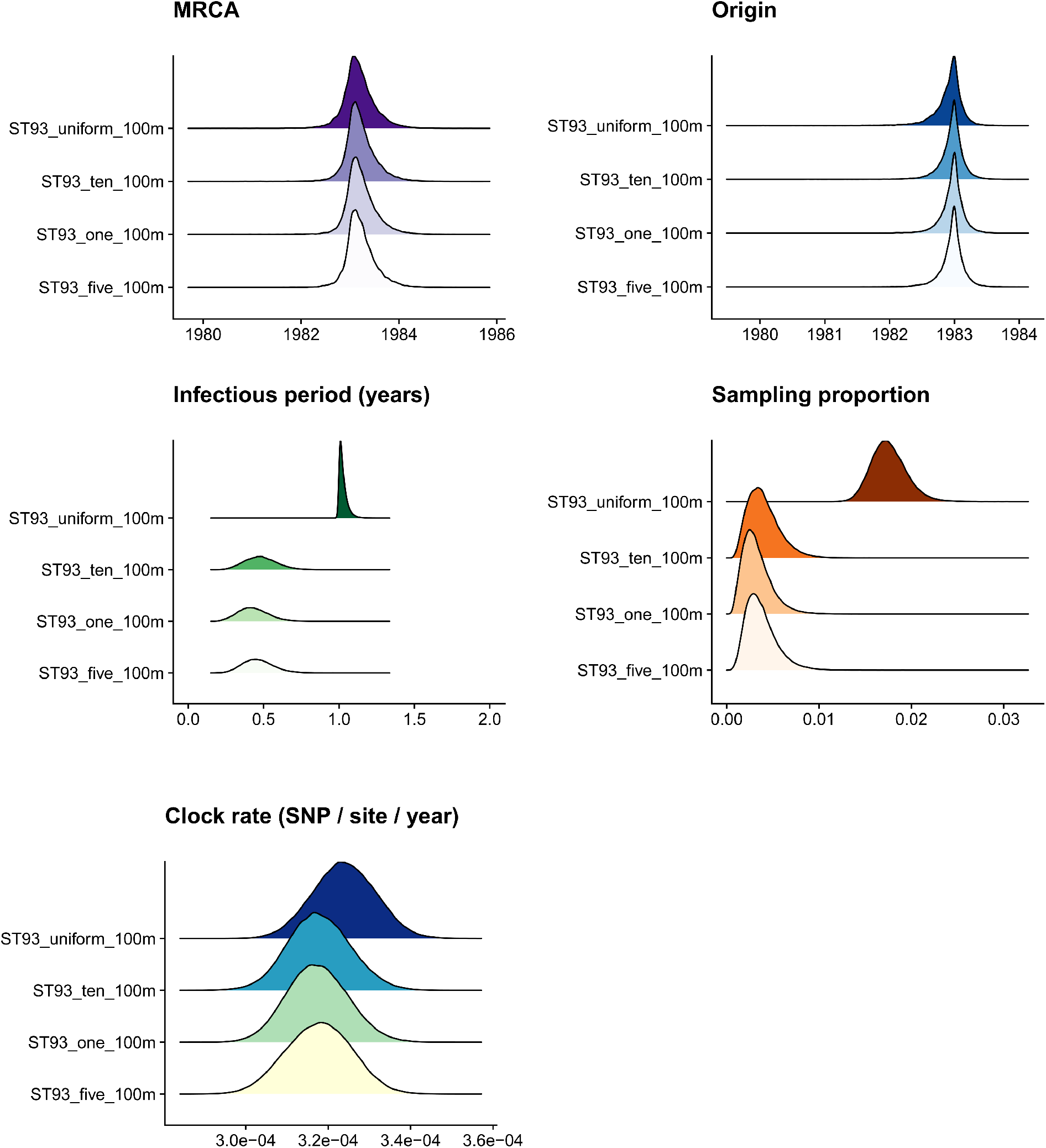
Posterior estimates of prior exploration in ST93 showing on the ridges the become uninfectious rate posteriors, where the prior was configured as a flat *Beta* distribution, or with *Lognormal*(*µ*, 1.0) with *µ* = 0.1 (10 years infectious period), *µ* = 0.2 (5 years) and *µ* = 1.0 (1 year). Full data on other lineages available in data repository.

**Fig. S6.**
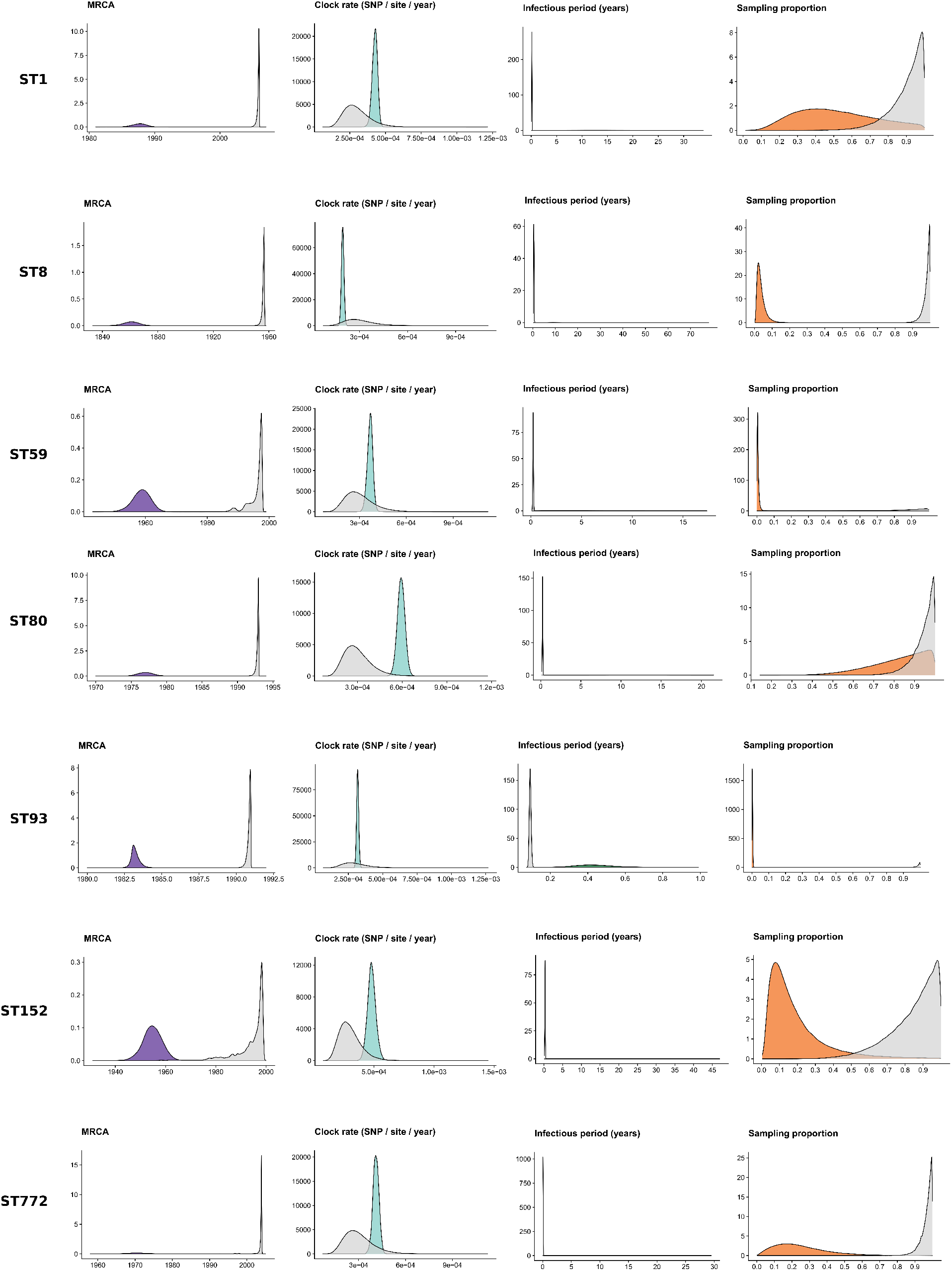
Prior sensitivity analysis of the birth-death skyline model for main sequence types of *Staphyloccocus aureus* (Table 1), showing the posterior distributions of the MRCA, clock rate, infectious period (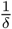) and sampling proportion (*ρ*) (colors), and the posterior distributions ‘under-the-prior’ (gray, without including the sequence alignment; necessarily including dates that can still inform the posterior estimates). Full data available in data repository.

**Fig. S7.**
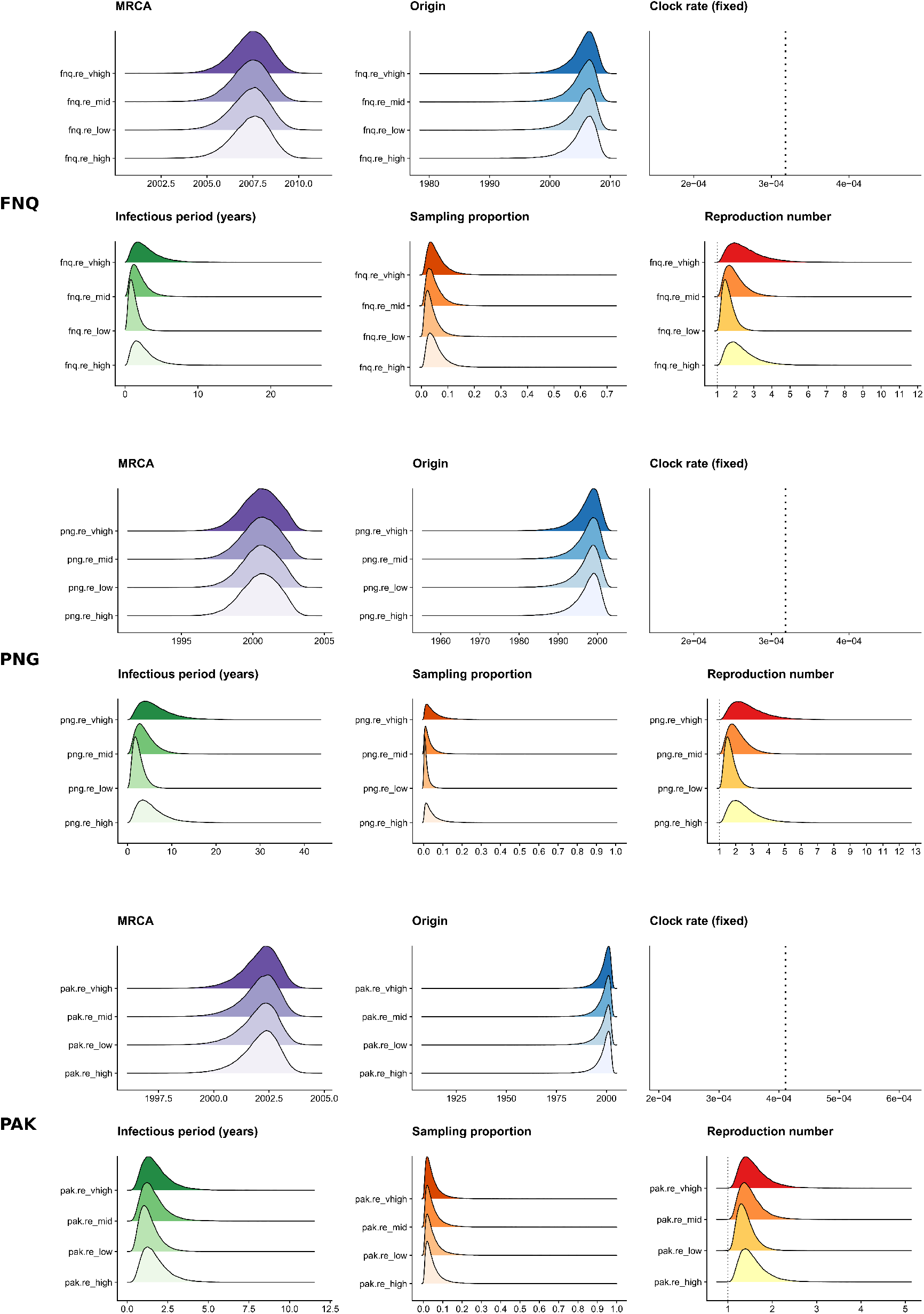
Birth-death skyline posteriors of three outbreaks sequenced in this study (ST93-MRSA-FNQ, ST93-MRSA-PNG, ST772-MRSA-PAK) across four configurations of the R_e_ prior with *Gamma*(2.0, *θ*) where labels in the plots correspond to *θ* = 0.5 (re_low), *θ* = 1.0 (re_mid), *θ* = 1.5 (re_high), *θ* = 2.0 (re_vhigh).

**Fig. S8.**
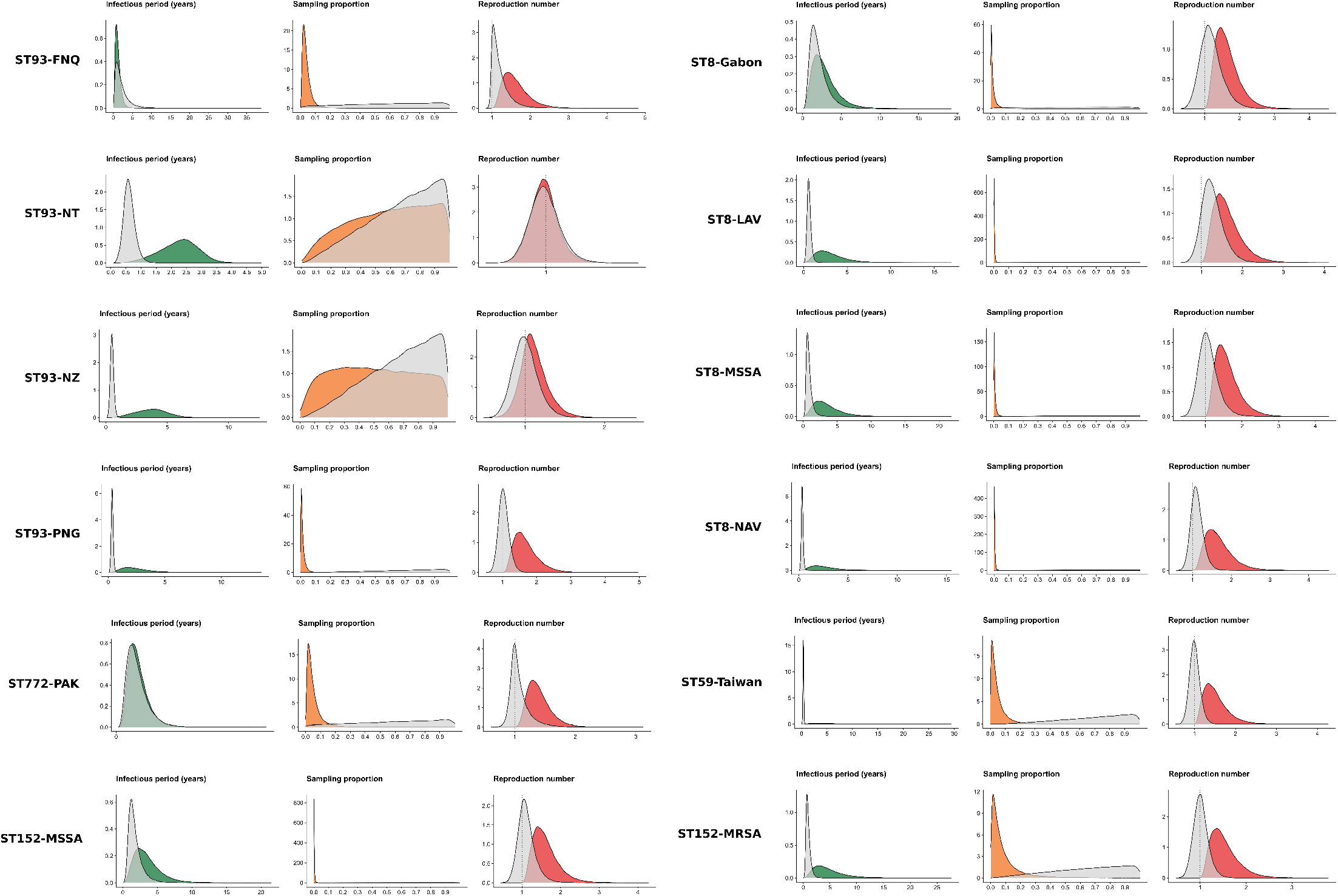
Prior sensitivity analysis of the birth-death skyline model for sublineages of *Staphyloccocus aureus* (Table 1), showing the posterior distributions of the infectious period 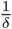, sampling proportion (*ρ*) and reproduction number (R_e_) (colors) and the posterior distributions ‘under-the-prior’ (gray, without including the sequence alignment; necessarily including dates that can still inform the posterior estimates). Full data available in data repository.

